# Epigenetic and genetic population structure is coupled in a marine invertebrate

**DOI:** 10.1101/2022.03.23.485415

**Authors:** Katherine Silliman, Laura H. Spencer, Samuel J. White, Steven B. Roberts

**Affiliations:** Marine Resources Research Institute, South Carolina Department of Natural Resources, Charleston, SC 29412, USA; School of Aquatic and Fishery Sciences, University of Washington, Seattle, WA, USA

**Keywords:** oyster, DNA methylation, single nucleotide polymorphism, Ostrea, environment, epigenetic

## Abstract

Delineating the relative influence of genotype and the environment on DNA methylation is critical for characterizing the spectrum of organism fitness as driven by adaptation and phenotypic plasticity. In this study, we integrated genomic and DNA methylation data for two distinct Olympia oyster (*Ostrea lurida*) populations while controlling for within-generation environmental influences. In addition to providing the first characterization of genome-wide DNA methylation patterns in the oyster genus *Ostrea*, we identified 3,963 differentially methylated loci between populations. Our results show a clear coupling between genetic and epigenetic patterns of variation, with 27% of variation in inter-individual methylation differences explained by genotype. Underlying this association are both direct genetic changes in CpGs (CpG-SNPs) and genetic variation with indirect influence on methylation (mQTLs). The association between genetic and epigenetic patterns breaks down when comparing measures of population divergence at specific genomic regions, which has implications for the methods used to study epigenetic and genetic coupling in marine invertebrates.

**Significance statement:** We know that genotype and epigenetic patterns are primarily responsible for phenotype, yet there is a lack of understanding to what degree the two are linked. Here we characterized the mechanisms and the degree by which genetic variation and DNA methylation variation are coupled in a marine invertebrate, with almost a third of the methylation variation attributable to genotype. This study provides a framework for future studies in environmental epigenetics to take genetic variation into account when teasing apart the drivers of phenotypic variation. By identifying methylation variation that cannot be attributed to genotype or environmental changes during development, our results also highlight the need for future research to characterize molecular mechanisms adjacent to genetic adaptation for producing long-term shifts in phenotype.

## Introduction

It is increasingly evident that epigenetic processes both influence phenotype and interact with genetic variation. One such epigenetic process is DNA methylation, which commonly refers to the methylation of a cytosine in a CpG dinucleotide. The role of DNA methylation is diverse across taxa and varies based on the genomic location. In most vertebrates, DNA methylation is widespread across the genome and silences transcriptional activity when present in the promoter regions (Wagner et al. 2014; Zemach et al. 2010; Varriale 2014). In contrast many marine invertebrates have sparsely methylated genomes and the influence of methylation on transcription is more complex (Suzuki & Bird 2008; Roberts & Gavery 2012; de Mendoza et al. 2019). In both vertebrates and invertebrates, the removal and addition of methyl groups can become canalized during the lifetime of that organism, and if occurring in germ cells has the potential to influence subsequent generations. This heritability of DNA methylation, as well as taxa-specific methylation rates and patterns, suggest that methylation differences arose in part due to evolutionary forces (Varriale 2014).

While the patterns and functions of CpG methylation differ among vertebrate and invertebrate taxa, in both systems methylation is highly variable. The evolutionary source of this variation is now an area of active research, with the two dominant factors appearing to be 1) an organism’s environmental history (intra- and inter-generational), and 2) its genotype (Jaenisch & Bird 2003; Lienert et al. 2011; Danchin et al. 2011). Understanding how the environment and genotype interact to influence DNA methylation is critical for delineating organism fitness as driven by phenotypic plasticity and adaptation, particularly in the context of global climate change. Bivalves, and oysters in particular, are a valuable model for investigating invertebrate methylation patterns due to their experimental tractability and concordant development of genomic resources (Timmins-Schiffman et al. 2013).

DNA methylation has been shown to vary in response to environmental factors in marine invertebrates (Eirin-Lopez & Putnam 2019). In oysters, differential methylation has been reported in response to ocean acidification (Lim et al. 2020; Downey-Wall et al. 2020), salinity stress (Xin Zhang et al. 2017), air exposure (X. Zhang et al. 2017), and the herbicide diuron (Akcha et al. 2020). Because there are clear associations between methylation and transcriptional activity (Gavery & Roberts 2013; Olson & Roberts 2014; Rivière 2014; Johnson et al. 2020; Song et al. 2017), methylation changes may contribute to phenotypic plasticity in response to abiotic stressors (Venkataraman et al. 2020; Wang et al. 2021a; Lim et al. 2020; Gonzalez-Romero et al. 2017; Wang et al. 2020; Downey-Wall et al. 2020). Methylation changes triggered by the environment may themselves be heritable if they occur in gametes, leading to transgenerational plasticity. It is the dynamic characteristics of the methylome that is fueling a growing body of research associating methylation variation with environmental exposures, particularly in trans-generational studies (Eirin-Lopez & Putnam 2019). However, few studies have controlled for (or described) the relationship between methylotype and genotype in test organisms (but see (Wang et al. 2021b; Johnson & Kelly 2020; Kvist et al. 2018), likely because in non-model taxa there is limited understanding of how the methylome is shaped by the genome.

While efforts to explore the influence of genotype on DNA methylation are limited in marine invertebrates, studies in taxa with advanced genomic resources have identified associations between genetic variants and DNA methylation (Banovich et al. 2014; Taudt et al. 2016). In oysters, genes with constitutive high levels of methylation have less genetic variation within populations (Roberts & Gavery 2012). Similar results have been found in the coral *Apis mellifera* and the jewel wasp (*Nasonia vitripennis*)(Lyko et al. 2010)(Park et al. 2011). One direct mechanism by which genetic and epigenetic variation can be associated are single nucleotide polymorphisms (SNPs) that create or remove CpG loci (CpG-SNPs), and therefore can immediately affect local methylation status (Shoemaker et al. 2010; Zhi et al. 2013). Alternatively, methylation status itself may change the likelihood of a SNP from occurring by “shielding” genetic mutations from selection, allowing genetic differentiation to accumulate (Klironomos et al. 2013), and by changing rates of homologous recombination (Li et al. 2012) and copy number variation mutation (discussed in (Skinner et al. 2014)). Surprisingly, some recent studies in oysters have found no relationship between genetic and epigenetic differentiation among populations or breeding cohorts, resulting in the suggestion that these two processes are uncoupled (Johnson & Kelly 2020; Jiang et al. 2013; Wang et al. 2020).

Genetic changes that are associated with methylation state but located some distance from the associated CpG are referred to as methylation quantitative trait loci (mQTLs). In humans, mQTLs may contribute up to 15-20% of inter-individual variation in methylation and up to 70% of population-level methylation variation (Heyn et al. 2013; McClay et al. 2015; Husquin et al. 2018; van Dongen et al. 2016). These genetic and epigenetic variants are often associated with complex traits or environmental differences, such as immunity or history of tobacco exposure (Gao et al. 2017; Bonder et al. 2017; McClay et al. 2015). Mechanistically, mQTLs have been proposed to operate in a number of ways. Global methylation patterns can be influenced by changing the expression or activity of methyltransferases, although mQTLs are rarely found in these genes. Increasingly, transcription factors and their binding sites have been implicated with mQTLs, as transcription factor binding can prevent methylation of nearby CpGs (Héberlé & Bardet 2019). Under this model, genetic variants in transcription factor binding sites can influence local methylation (local mQTLs), while genetic variants that affect the activity of wide-acting transcription factors can influence methylation at many distant CpGs near binding sites for that specific transcription factors (distant mQTLs). While these mechanisms have not been investigated in most non-model taxa, the conserved roles of transcription factors across taxa suggests that they may also play a role in shaping methylation variation in invertebrates and bivalves (Nitta et al. 2015; Bell et al. 2011). Functional genomics are needed to further investigate these relationships to ascertain the mechanisms underlying genetic and epigenetic relationships in nonmodel taxa.

The Olympia oyster (*Ostrea lurida*) is an emerging model taxa for investigating the links between environment, genetic adaptation, and epigenetic plasticity (White et al. 2017; Silliman 2019; Maynard et al. 2018; Timmins-Schiffman et al. 2013). Native to estuaries from Baja California to the central coast of Canada, *O. lurida* extends over strong environmental clines and mosaics (Chan et al. 2017; Schoch et al. 2006). Significant neutral and putatively adaptive genetic variation has been detected between populations at both regional and local scales, which is surprising given the potential for high connectivity during the planktonic larval phase (Silliman 2019). Experimental tests for local adaptation among neighboring sites within San Francisco Bay, CA (Maynard et al. 2018) and Puget Sound, WA (Silliman et al. 2018; Heare et al. 2017) have found phenotypic variation at fitness-related traits, such as growth, salinity tolerance, and reproductive timing. By controlling for environmental variation, these studies suggest a strong heritable component underlying population differences. Whether this component is due to genetic variation, inherited epigenetic modifications, or a combination is still unknown.

The objective of the current study was to leverage a new *O. lurida* draft genome to investigate the relationship between CpG methylation and genetic variation based on 2b-RAD SNPs. Oysters from two populations in Puget Sound, WA were raised to maturity for one generation in common conditions to remove lifetime exposure to environmental variation. These populations have phenotypic variation in larval and adult size and reproductive timing (Silliman et al. 2018; Heare et al. 2017; Spencer et al. 2020), show varied gene expression profiles under stress (Heare et al. 2018), and come from sites with different environmental profiles in dissolved oxygen, temperature, pH, salinity, and food availability (Moore et al. 2008; Banas et al. 2015; Khangaonkar et al. 2018). While some marine invertebrate studies have associated overall patterns of epigenetic and genetic differentiation between populations (e.g., (Johnson & Kelly 2020), to our knowledge this is the first to directly link epigenetic and genetic variability by identifying and functionally characterizing CpG-SNPs and meth-QTLs.

## Results

### Study Design

Adult Olympia oysters (*O. lurida*) derived from two separate parent populations in Puget Sound, Washington were reared in Clam Bay, Washington. The two parental populations were from Hood Canal and South Sound. Details on broodstock collection and outplanting are described in (Heare et al. 2017). Shell length and wet weight were measured immediately prior to adductor tissue dissection. Biallelic SNPs were genotyped in 114 individuals (57 from each population) using a reduced-representation 2b-RAD approach (Wang et al. 2012) by mapping to a draft *O. lurida* genome assembly (GCA_903981925.1) (159,429 scaffolds, N50 = 12,947). After filtering for sample coverage (at least 3 reads in >70% of individuals) and a minimum overall minor allele frequency (MAF) of 0.01, genotype likelihoods were calculated with ANGSD for 5,269 SNPs and used for subsequent population genetic analyses (Korneliussen et al. 2014).

To characterize CpG methylation patterns, we randomly selected 9 genotyped individuals from each population and used methyl-CpG binding domain (MBD) bisulfite sequencing (MBD-BS). These 18 samples are referred to as the MBD18 samples. This reduced representation approach is efficient for taxa with sparse methylation patterns, as it enriches for methylated DNA regions while providing single base resolution through bisulfite conversion (Trigg et al. 2021). Reads from all MBD18 samples were concatenated into one ‘meta-sample’ and aligned to the *O. lurida* genome to describe general methylation patterns in the Olympia oyster. Out of 2,030,624 CpG loci with at least 5x sequencing coverage in the ‘meta-sample’, 1,839,241 (90.6%) were methylated, defined as loci with greater than 50% of reads remaining cytosines after bisulfite conversion.

For comparative methylation analyses, reads from each MBD18 sample were aligned separately, and a more conservative set of 252,115 loci were used by filtering for loci with 5x coverage across at least 7 of the 9 samples within each population. As MBD-BS enriches for methylated regions, this conservative filtering approach may exclude regions that were methylated in one population but largely unmethylated in the other. Therefore, we included an additional 251 CpG loci that were minimally sequenced in one population (≤1 sample) and widely sequenced in the other population (≥7 samples), and annotated samples with missing data in the low-sequenced population as unmethylated at 5x coverage.

### Genome annotation and general methylation landscape

The draft genome assembly (Accession # GCA_903981925.1) is 1.1 Gb in size with a contig N50 of 7.8kb. Gene prediction identified 32,210 genes, 170,394 exons, and 163,637 coding sequences. Additionally, 27,331,887 CpG motifs were identified in the genome assembly.

Transposable element identification determined GC content of the genome to be 36.58%. Retroelements comprised 6.24% of the genome assembly. Those retroelements consisted of 0.03% small interspersed nuclear elements (SINEs), 5.69% long interspersed nuclear elements (LINEs), and 0.53% of long terminal repeat (LTR) elements. DNA transposons made up 3.13% of genome assembly.

Of the 27,331,887 CpGs in the *O. lurida* genome we found that 1,839,241 were methylated (6.73%) using the concatenated MBD18 reads. Of the 1,839,241 methylated loci 34.5% were intragenic (14.7% in exons, 19.8% in introns), 4.6% and 4.7% were located 2kb upstream and downstream of known genes, respectively, and 13.8% were within transposable elements. 32.3% of the methylated loci were not associated with known regions (i.e. intergenic beyond 2kb gene flanking regions) (Figure 1). The distribution of methylated loci across genomic features differed significantly from the distribution of all CpG loci in the *O. lurida* genome (*χ*^2^=685,890, df=5, p∼0), and methylated CpG loci were ∼3.7x more likely to be located within an exon (Supplemental Figure 2).

**Figure 1:**
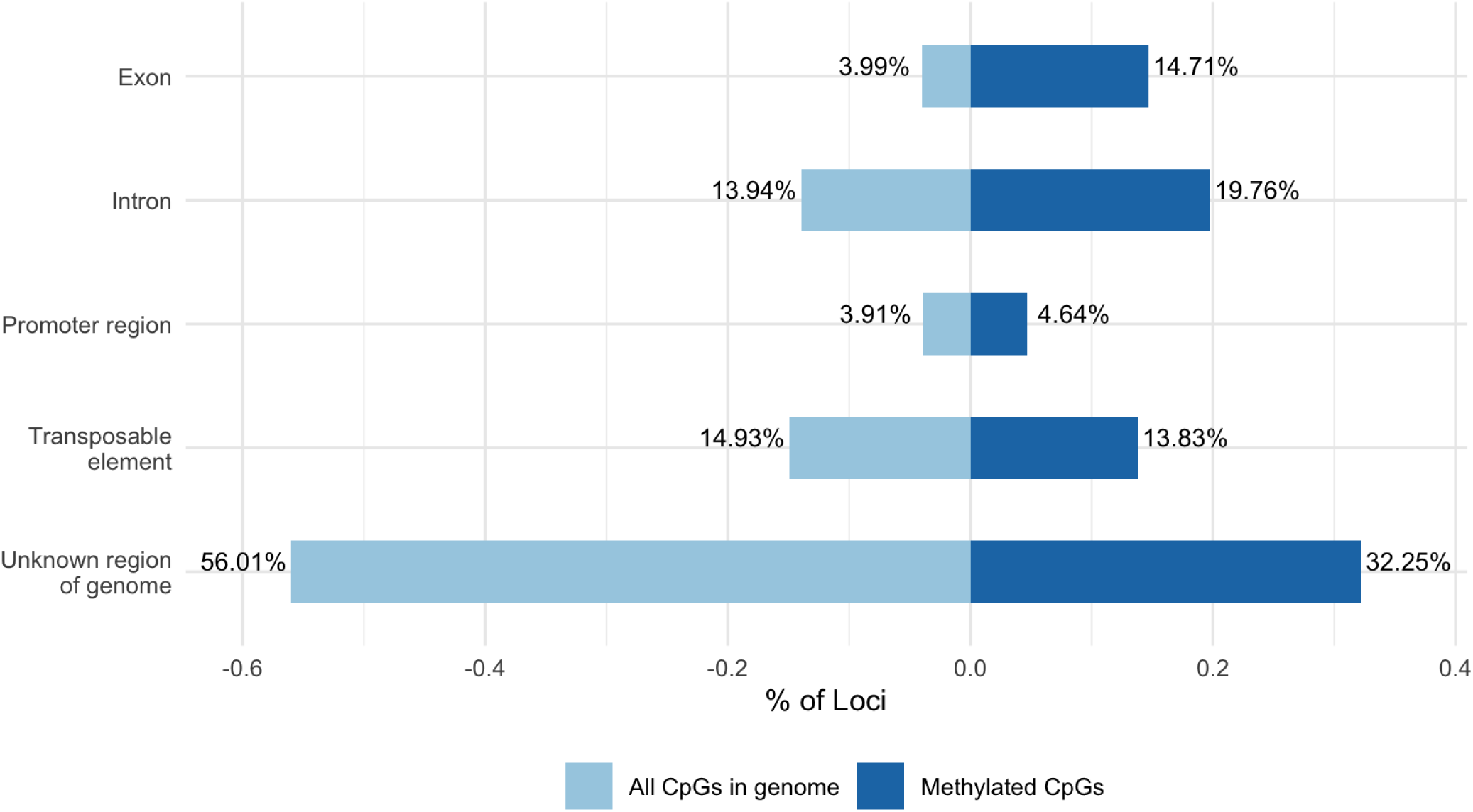
Comparison of the percentages of CpGs in genome (light blue) vs. methylated CpG loci (dark blue) in *O. lurida* muscle tissue that intersect with each of the following genomic features: exon, intron, promoter region (within 2kb of the 5’ end of a gene), transposable element, unknown region of genome. Compared to all CpGs in the *O. lurida* genome methylated loci are more likely to be located in exons (3.7x) and introns (1.4x), and less likely to be located in unknown regions (0.60x).

### Population genetic structure

Population genetic analyses of all 114 individuals found evidence of divergence with gene flow between the two populations. Principal component analysis (PCA) of 5,269 SNPs clustered individuals primarily by population of origin along PC1, which represented 6.64% of the total variation (Figure 2). NGSadmix was used to perform an ADMIXTURE analysis based on genotype likelihoods of 3,724 SNPs, after filtering further for a minimum overall allele frequency of 0.05 (Skotte et al. 2013). The most likely number of genetic clusters (K) was determined to be K = 2, with evidence of admixture between the two sampled populations (Supplemental Figure 7). Outlier analyses with BayeScan detected 12 SNPs as potentially under divergent selection (Foll & Gaggiotti 2008). One of these SNPs was found in a gene involved in cell mitosis (G2/mitotic-specific cyclin-B) and another was within 2kb downstream of a gene involved in protein ubiquitination (SOCS5) (Supplemental Table 3).

**Figure 2:**
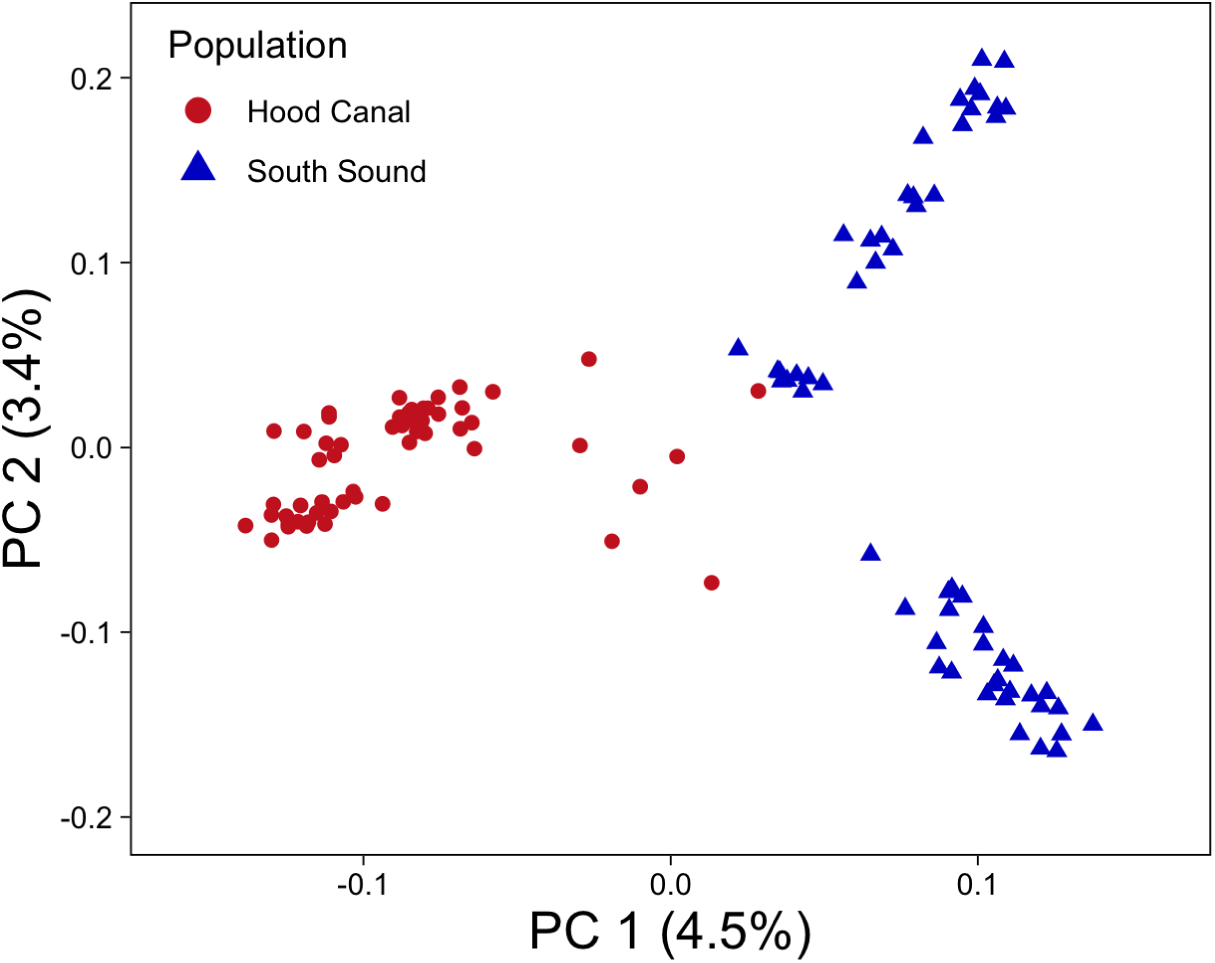
PCA based on 5,269 SNPs for 114 individuals, with colors and shape referring to parental population.

Population genetic differentiation (F_ST_) was measured overall and separately for each SNP and each gene region (± flanking 2 kb) (Reynolds et al. 1983). These values were derived from estimates of the site-frequency spectrum (SFS) in ANGSD, and therefore used 5,882 SNPs that were filtered as to avoid distorting the allele frequency spectrum (Korneliussen et al. 2014). Overall unweighted F_ST_ between the two populations was 0.0596 (SD=0.087), and weighted Fst was 0.0971. Per-SNP F_ST_ was calculated for 5,882 SNPs, with 1,909 of these SNPs found across 1,386 genes. Mean F_ST_ for SNPs in genes was slightly lower than overall F_ST_ with an unweighted F_ST_ of 0.0586 (SD=0.084) and weighted F_ST_ of 0.093. 38 genes had an F_ST_ > 0.3, and were enriched for four biological processes, including steroid hormone mediated signaling pathway and three processes related to autophagy (Supplemental Table 4).

### DNA methylation differences between populations

Population methylation analyses of the MBD18 individuals found evidence of epigenetic divergence. Principal component analysis, which was performed on a percent methylation matrix (252,366 loci x 18 samples), clustered individuals by population of origin along PC2, which represented 8.5% of the total variation (Figure 3b). Logistic regression analysis identified 3,963 differentially methylated loci between populations (DMLs, methylation difference >25% and Q-value <0.01, Supplemental Figure 4), 1,915 of which were located within known genes (48.3% of DMLs), and 1,504 of which were within exons (40.0% of DMLs). An additional 178 and 171 of the DMLs were found upstream and downstream of genes (within 2kb; 4.5% and 4.4%, respectively), and 188 were located within transposable elements (4.7%). There were 500 DMLs that were not found in any known feature (12.6%). 54% of DMLs had higher methylation levels in SS (2,154 loci), and 46% were higher in HC (1,809) (Supplemental Figure 2).

**Figure 3:**
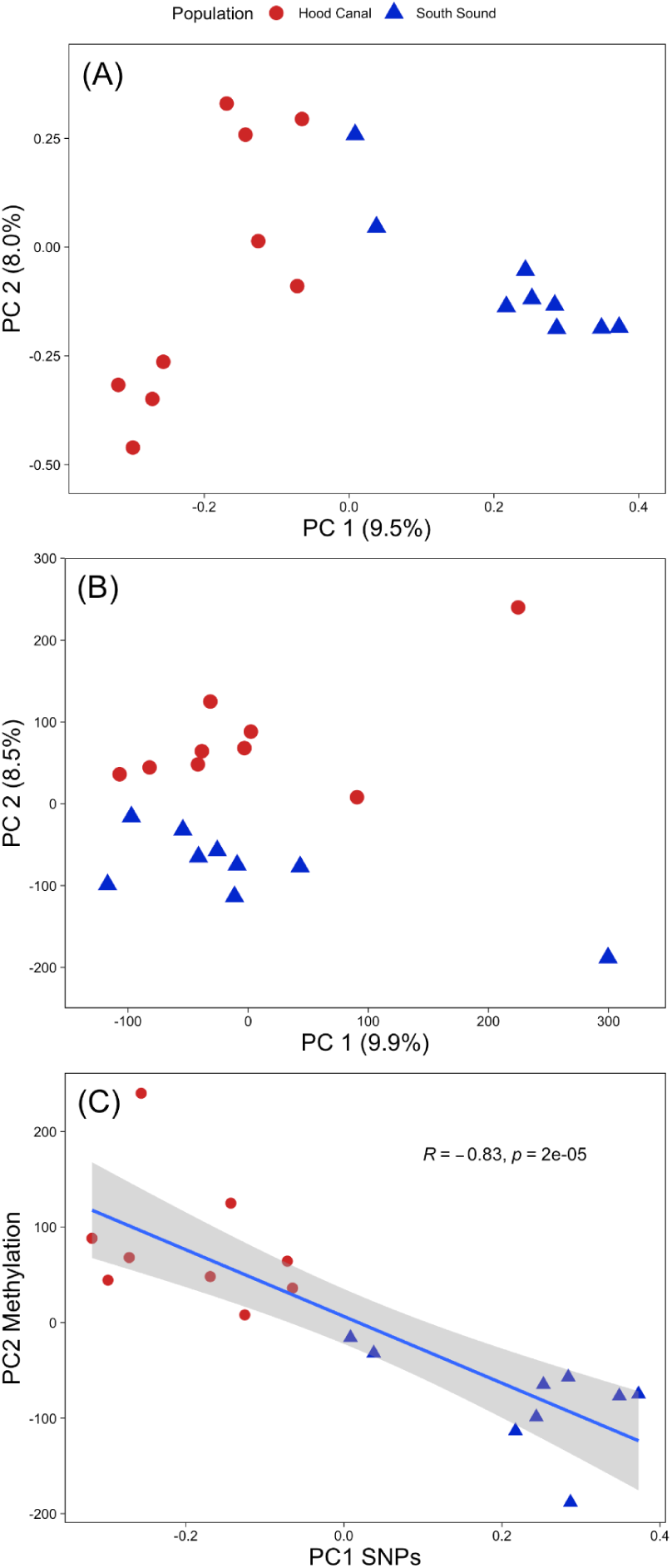
**A.** PCA of SNP data for the MBD18 samples. **B.** PCA of DNA methylation data (using all loci) for MBD18 samples. **C.** Scatter plot of PC1 from SNP genotype data and PC2 from DNA methylation data showing the linear regression line, Pearson correlation coefficient, and p-value.

Population divergence of methylation was also assessed at the gene level for gene regions containing ≥5 informative loci. Of the 6,299 gene regions assessed, 1,447 were differentially methylated (DMGs) as determined by binomial GLMs. DMGs and gene regions containing DMLs were each enriched for 31 biological processes, both of which included sarcomere organization (GO:0045214), and metabolic process (GO:0008152) (Supplemental Table 2). Mean P_ST_, a measure of population divergence in methylation (Johnson & Kelly 2020), averaged across 14,088 random 10kb bins was 0.30 ±SD 0.26.

### Relationship between genetic and epigenetic variation

To investigate the relationship between genetic and DNA methylation variation, we first compared pairwise genetic distances between MBD18 samples with pairwise Manhattan distance based on all filtered methylation data and found a weak, but significant relationship (Pearson’s *R* = 0.27, p-value =0.00077 and Spearman’s ρ = 0.22, p-value = 0.0069, Figure 4a). This correlation was stronger when comparing against Manhattan distances based on DMLs (Pearson’s *R* = 0.73, p-value <2.2E10^-16^ and Spearman’s ρ = 0.70 p-value < 2.2E10^-16^) (Figure 4b). This suggests that the rate of genetic changes between individuals is similar to the rate of methylation changes, especially for CpG sites that diverge between populations. We further compared population specificity of our data by correlating the 1st PC scores from SNP data (9.5% of variation, Figure 3a) with the 2nd PC scores of methylation data (8.5%, Figure 3b), as these two axes clearly separated individuals by population. These were strongly correlated (Pearson’s *R* = -0.83, p-value = 1.65E10^-5^ and Spearman’s ρ = -0.86, p-value < 2.2E10^-16^, Figure 3c), suggesting that common underlying mechanisms may be involved in population divergence at variable genetic and epigenetic sites. However, we found no significant correlation between F_ST_ and P_ST_ at 827 random 10kb genomic bins where we had both SNP and methylation data (Figure 5). This result suggests that the strong correlation between population-specific genetic and epigenetic patterns on the individual level is not primarily driven by genomically linked epigenetic and genetic sites.

**Figure 4:**
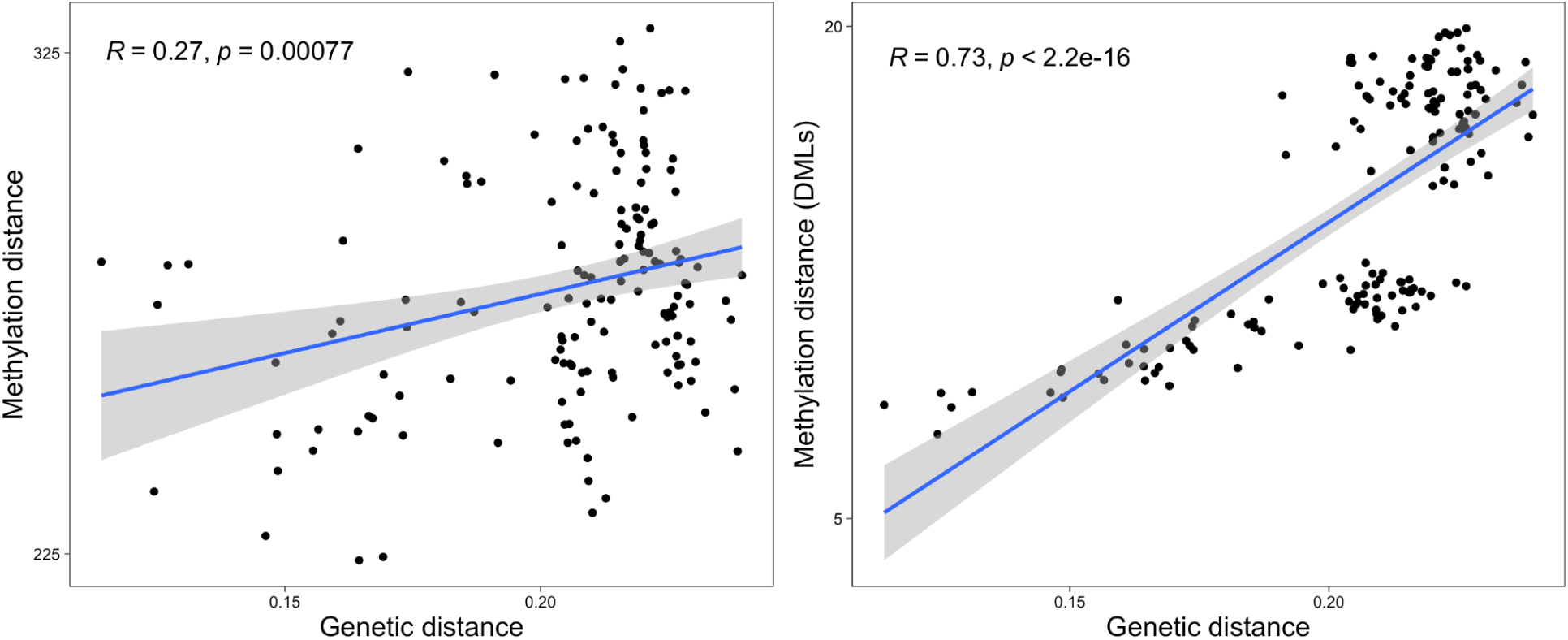
Epigenetic divergence as a linear function of genetic distance. The x axes represent genetic distances calculated from genotype probabilities for 5,269 SNPs. The y axes are the Manhattan distances from CpG methylation x1000 (a; using all methylation data and b; using DMLs). The linear regression lines are shown, together with the Pearson correlation coefficient and p-value.

**Figure 5:**
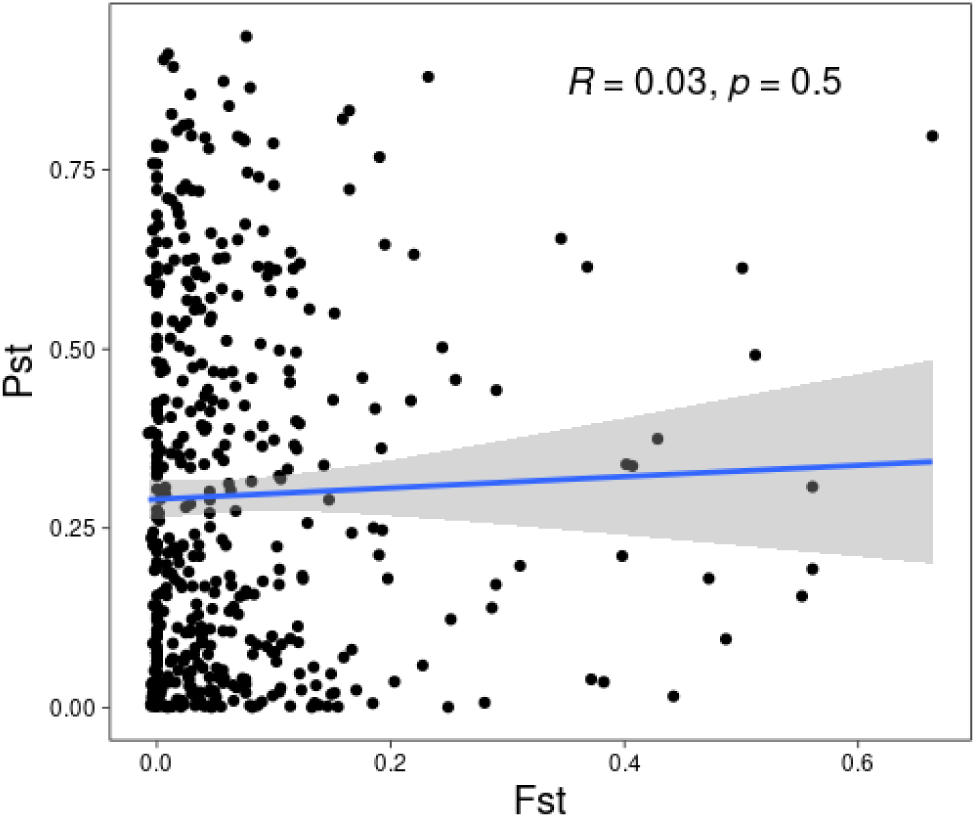
Scatterplot of P_ST_ (measure of epigenetic divergence between populations) and F_ST_ (measure of genetic divergence) for 827 random 10kb genomic bins with both SNP and methylation data.

### mQTL analysis

To determine if genetic variants are associated with loci showing inter-individual methylation variation, we conducted a mQTL analysis using a linear regression model in the R package MatrixEQTL (Shabalin 2012). For this analysis, we used 2,860 SNPs that had a MAF > 0.05 across the MBD18 samples as the explanatory variable, PC1-3 of SNP genotype data as a covariate to control for ancestry, and the percent methylation at 232,567 CpG sites as the response variable. ‘Local’ mQTLs were determined to be SNPs within 50kb of the CpG and an un-adjusted p-value threshold of 0.01, while distant mQTLs were greater than 50kb from the CpG or on a different scaffold and had an FDR threshold of 0.05. Results of the mQTL analysis are summarized in Table 1, with 1,985 SNPs (69.4%) detected as mQTLs and 7,157 CpGs (3.1%) associated with a mQTL. Due to linkage disequilibrium (LD) among SNPs as well as our reduced representation genetic sequencing, most of these SNPs are unlikely to be the actual causal variant influencing the methylated site. Therefore, we follow the recommendation of (McClay et al. 2015) and evaluate the methylated sites under genetic control as a better representation of genetic influence on the methylome.

**Table 1.** Summary statistics for mQTL analyses.

|  | Local (< 50kb)<br>p < 0.01 | Distant (> 50kb,<br>different scaffolds)<br>FDR < 0.05 |
| --- | --- | --- |
| Number of CpG loci tested | 10,320 | 232,567 |
| Number of SNPs tested | 853 | 2,860 |
| Number and percent of unique SNPs with mQTLs | 181 (21.2%) | 1,936 (67.7%) |
| Number and percent of unique methylated sites with mQTLs | 240 (2.3%) | 6,926 (3.0%) |
| Number and percent of methylated sites associated with an mQTL<br>and a CpG-SNP / Number of tested sites with a CpG-SNP | 12/179 (6.7%) | 8/273 (2.9%) |
| Number and percent of methylated sites with an mQTL and a DML | 58/240 (24.1%) | 655/6926 (9.5%) |

Compared to background rates, local mQTLs were overrepresented in gene regions (83 annotated genes, 67% vs. 37%, *p*=8.665 E-16), as were their associated CpGs (78% vs. 59%, p=7.23E-9) (Supplemental Figure 9). Genes containing these sites were functionally enriched for the GO term “DNA repair” (8.7% of genes), InterPro term “SWI/SNF chromatin-remodeling complex” (3.6%), and UP keywords “transcription regulation” (16.8%) and “disease mutation” (18%), among other functions. Distant mQTL SNPs were found in 309 genes but were not enriched for any functional categories. The CpG loci associated with distant mQTLs were found in 1,809 annotated genes and enriched for the COG category “RNA processing and modification” (0.7% of genes), 49 GO categories including “transcription DNA-templated” (12.1%), “mRNA processing” (2.6%), “covalent chromatin modification” (2.4%), “regulation of translational initiation” (0.66%), “chromatin remodeling” (1.1%), nucleic acid binding (5.7%), chromatin binding (3.8%), and “transcription factor activity”(3.8%), as well as 87 UP keywords and sequence features including “phosphoprotein”(49.7%), “nucleus” (33.2%), “acetylation” (22%), “RNA-binding” (6.7%), “methylation” (6.4%), and “chromatin regulator” (3.9%). Some genes containing these distantly controlled CpGs include 7 different RNA binding motif proteins, 6 RNA polymerase genes, 8 DEAD-box type helicases, 17 eukaryotic translation initiation factor (eif) genes, and six SWI/SNF regulator of chromatin. While most other enrichment tests presented here were not significant after Benjamini FDR correction (P < 0.1), 17 (10%) of the enriched functions for CpGs with distant mQTLs were significant (Supplementary File 2).

SNPs that create or remove CpGs (CpG-SNPs) may contribute to individual differences in methylation, and therefore lead to mQTL associations or correlations between genetic and epigenetic distances. We identified 651 CpG-SNPs (12.4%) from our full set of 5,269 SNPs through mapping to our draft genome. CpG-SNPs were more likely to be within 350bp of a methylated CpG than non-CpG-SNPs (40.9% vs 34.5%, p=0.00161), when using all CpGs in the genome as the background. This 350bp window size represents the maximum length of a library fragment, and therefore is the maximum distance at which a CpG-SNP could directly affect our measure of methylation. CpG-SNPs were slightly more likely to be within 350bp of a DML than non-CpG-SNPs (0.9% vs 0.4%, p=0.04086), suggesting that CpG-SNPs only play a minor role in creating DMLs. More of the methylated sites with local meth-QTLs had a CpG-SNP compared to distant mQTLs (12 vs 8, p=1.508e-13). Due to the sparse genetic sequencing of the genome, we are likely missing many CpG-SNPs associated with both local and distant mQTLs.

We investigated the spatial overlap between population DMLs and CpGs associated with local mQTLs, and found CpGs with local mQTLs were more likely to overlap with DMLs than CpGs without an mQTL association (24.1% vs. 15.9%, p=0.001289). Genes of particular interest included PRICKLE2 (developmental processes, linked to growth in *Crassostrea gigas (Takeuchi et al. 2003; Yang et al. 2020)*, TRIM2 (innate immunity, differential methylation to low pH in *C. hongkongensis* larvae (Ozato et al. 2008; Lim et al. 2020)), eukaryotic translation initiation factor 3 (eif3) (translation initiation through mRNA recruitment and interactions with methyltransferases, response to low pH in *Saccostrea glomerata* (Wolf et al. 2020; Ertl et al. 2016), OXCT1 (ketone catabolic process, variably methylated in humans (Feng et al. 2021)), Mapk6 (signal transduction, immune signaling in *S. glomerata (Ertl et al. 2016)*), and MLH3 (DNA mismatch repair protein (Lipkin et al. 2000)). Examples of two local mQTLs that are also a DML are shown in Figure 6. A significantly lower proportion of distant mQTLs were associated with DMLs (9.5%, p = 1.243e-8). Of these 655 sites, 363 were in 311 genes, which were enriched for numerous processes relative to all distant mQTL genes, including functions related to development, immune response, transcription factor activity, and coiled coil domains. Unlike local mQTLs, distant mQTLs were deficient in DMLs relative to non-distant mQTLs (9.5% vs 14%, p < 2.2 10e-16).

**Figure 6:**
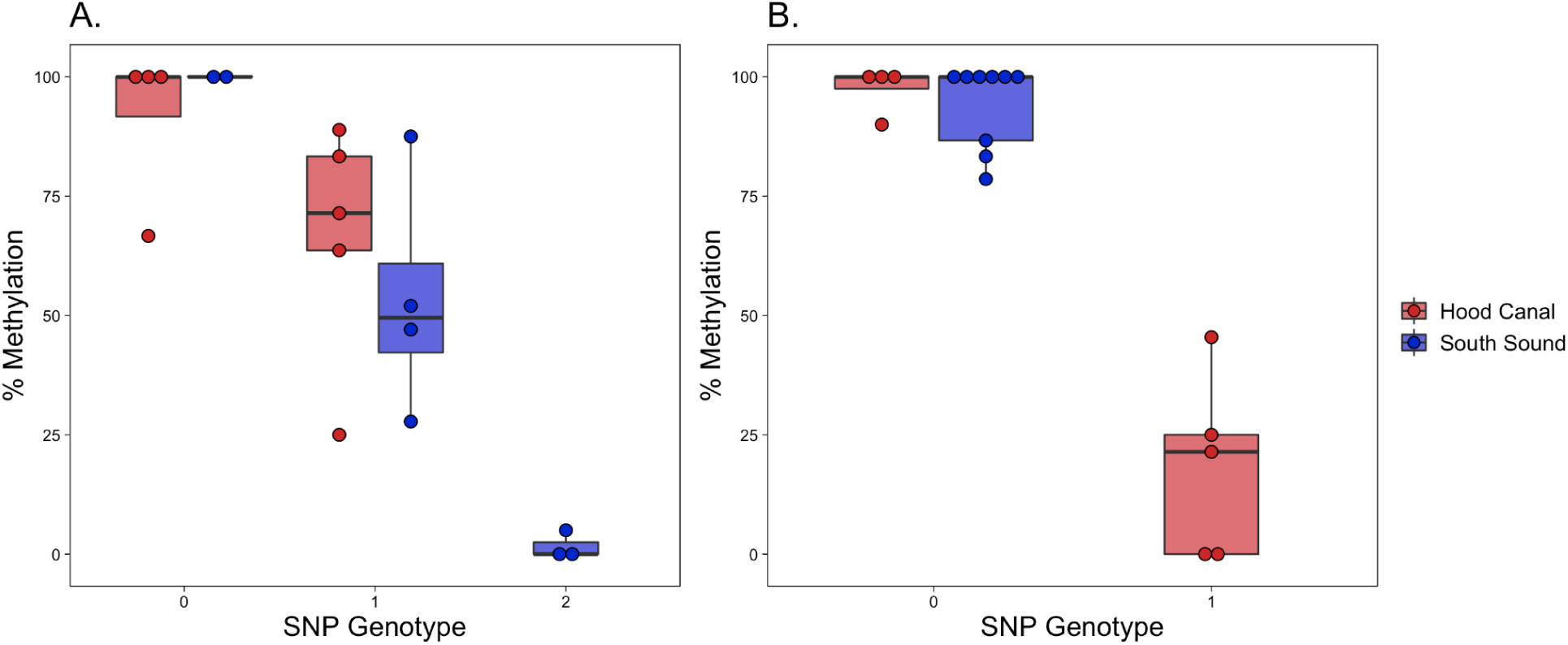
Two example CpG loci that are associated with a local SNP and are differentially methylated between populations. Each dot is an individual, with the genotype of the SNP on the x-axis and percent methylation on the y-axis. Boxplots are grouped and colored by population. A) CpG (Contig54624.19738) is found on a gene annotated as “similar to eif3d”, is differentially methylated 38.6% between populations, and associated with SNP Contig54624.19920. B) CpG (Contig60108.5780) is found on a gene annotated as “similar to MLH3”, is differentially methylated 37.3% between populations, and is associated with SNP Contig60108.2787.

## Discussion

Research primarily from humans and plants have shown that both environment and ancestry can influence variation in DNA methylation, however these associations are still not fully understood in less studied taxa such as marine invertebrates. In this study, we describe the genotype x epigenotype relationship by integrating high-throughput genomic and methylation data for two distinct oyster populations raised in the same environment for one generation. In addition to providing the first characterization of genome-wide methylation patterns in the oyster genus *Ostrea*, our results show a clear association between genetic and epigenetic patterns of variation. Underlying this association are both direct genetic changes in CpGs (CpG-SNPs), and mQTLs with indirect functional influence on methylation. The association between genetic and epigenetic patterns breaks down when comparing measures of population divergence at specific genomic regions, suggesting that individual variation can outweigh population-level variation when comparing these patterns at local genomic scales.

### General DNA methylation patterns

*O. lurida* CpG methylation is disproportionately found in gene bodies. When compared to all CpG loci in the genome, *O. lurida* methylation is ∼3.7x more likely to occur in exons, and ∼1.4x more likely to occur in introns (Figure 1, Supplemental Figure 2). Gene body methylation has also been reported for the Eastern oyster (*Crassostrea virginica*) (Venkataraman et al. 2020; Johnson & Kelly 2020; Downey-Wall et al. 2020), Pacific oyster (*C. gigas*) (Gavery & Roberts 2013; Song et al. 2017; Wang et al. 2020, 2014), Hong Kong oyster (*C. hongkongensis*) (Lim et al. 2020), and pearl oyster (*Pinctada fucata martensii*) (Zhang et al. 2020). The precise role and function of gene body methylation is not yet clear. However, in contrast to the suppressive role of promoter methylation in vertebrates, gene body methylation in invertebrates is hypothesized to mediate transcriptional activity because it is positively associated with gene expression (Roberts & Gavery 2012). Without expression data we cannot directly assess the relationship between genic methylation and transcription in *O. lurida.* However the high preponderance for methylation in *O. lurida* exons, and to a lesser extent introns, supports a role in mediating alternative splicing activity. That methylated genes in the *O. lurida* genome are enriched for a variety of biological processes, including those related to cell cycle and biogenesis, DNA, RNA and protein metabolism, transport, and stress response (Supplemental Table 1), supports the theory that methylation regulates both housekeeping and inducible processes in marine invertebrates.

### Epigenetic and genetic population structure

#### Population-specific methylation patterns

Global DNA methylation patterns in *O. lurida* are influenced by population of origin (Figure 3b), despite rearing oysters in common conditions. To examine biological functions associated with differential methylation among these populations, we performed enrichment analyses on both the 1,447 differentially methylated gene regions (DMGs) and genes containing the 3,963 differentially methylated loci (DMLs) (Supplemental Table 2). DMGs and genes containing DMLs were both enriched for biological processes involved in transport, cell adhesion and migration, protein ubiquitination, and sarcomere organization. DMGs were also enriched for 27 other processes, including several related to reproduction (e.g. germ cell development, lipid storage, oogenesis), and growth (e.g. cell morphogenesis, epithelium development, regulation of neurogenesis and growth). The two focal populations have distinct abiotic stress tolerances, as well as reproductive and growth strategies, some of which have been shown to be transgenerational (Silliman et al. 2018; Spencer et al. 2020; Heare et al. 2017). As gene expression is associated with methylation status in oysters (Gavery & Roberts 2013; Johnson et al. 2020), protein-coding genes identified here with population-specific methylation rates are good candidates for future studies exploring epigenetic control of phenotype in marine invertebrates.

Our methylation data is biased towards hyper-methylated loci (average proportion methylation for loci with 5x coverage is ∼80%, and only 2.5% of sequenced loci had no methylated reads)(Supplemental Figure 1). This type of data is excellent for characterizing the methylation landscape, but does limit our ability to compare loci where methylation varies significantly among populations (e.g. loci that are hyper-methylated in one population, but hypo-methylated in the other). To partially mitigate this concern, we implemented a filtering approach that is atypical of MBD-BS studies in order to include some such divergently methylated loci that may be missed with otherwise strict data filtering. Of these 251 included loci, 246 were DMLs and 17 were associated with distant mQTLs, supporting this choice when going forward with comparative MBDseq or MBD-BS. As the cost of sequencing decreases, other sequencing methods (e.g. WGBS) should be used to detect other regions where methylation differs substantially. However, by detecting population-specific epigenetic differences, our results contribute to the limited number of studies from *Crassostrea* oyster species that also found population-specific (Johnson & Kelly 2020; Zhang et al. 2018) or family-specific methylation patterns (Olson & Roberts 2014). In contrast to these previous studies, the present study controls for changes to the methylome that could arise due to differing environments during development. Therefore, the observed population-specific methylation patterns reflect either heritable methylation differences, or those acquired as germ cells in the parental environments.

#### Population genetic variation

Low but significant population genetic divergence had previously been described for Olympia oyster populations in Puget Sound using de novo genotype-by-sequencing and 2b-RAD data (Silliman et al. 2018; Silliman 2019). The current study validates these findings using a reference-based 2b-RAD approach and 5,269 SNPs, finding weak (F_ST_= 0.059), but significant genetic differentiation (Figure 2, Supplemental Figure 7). Similar genetic differentiation patterns are observed for other bivalve species on comparable spatial scales, such as the Eastern oyster (*C. virginica*) and the Pacific oyster (*C. gigas*) (Johnson & Kelly 2020; Kawamura et al. 2017). Given the potential for gene flow between neighboring oyster populations during the planktonic larval stage, the continued evidence for population genetic differentiation suggests that either larvae do not disperse as far as would be predicted (Shanks 2009; Pritchard et al. 2015), or that adaptive and neutral processes can override the effects of gene flow for some parts of the genome (Sanford & Kelly 2011; Weersing & Toonen 2009).

A benefit of using a reference genome in this study is the ability to also evaluate functional patterns of genetic divergence. SNPs in genes had lower mean F_ST_ than the genome-wide average, which aligns with expectations of gene bodies in general showing higher sequence conservation due to purifying selection (Kimura 1983). Two gene regions contained outlier SNPs, and therefore may be under divergent selection: G2/mitotic-specific cyclin-B and SOCS5. G2/mitotic-specific cyclin-B is associated with gametogenesis in *C. gigas (Dheilly et al. 2012)*, as well as tidally-influenced gene expression changes in the mussel *Mytilus californianus (Gracey et al. 2008)*. SOCS5, a member of the cytokine signaling family, is highly expressed in hemocytes, gills, and the digestive gland of *C. gigas* (Li et al. 2015). Our 2b-RAD SNPs only represented 1,386 genes out of 32,211 in the genome, and therefore our outliers are likely only a fraction of genes diverging between these populations (Lowry et al. 2017). Nevertheless, these genes should be added to a growing list of candidate loci to investigate further for local adaptation in the Olympia oyster (Silliman 2019; Heare et al. 2018; Maynard et al. 2018).

### Associations between methylation patterns and genetic variation

Previous studies associating genetic variation and DNA methylation patterns in marine invertebrates mainly compared measures of population divergence (e.g., F_ST_ and P_ST_) at overlapping genomic regions, and found little or no relationship (Johnson & Kelly 2020; Wang et al. 2020; Liew et al. 2020). In the current study, we also found no relationship between F_ST_ and P_ST_ for overlapping genomic regions. However, by further comparing genome-wide summary statistics and PCAs at the individual level, we revealed the significant relationship between interindividual patterns in methylation and genetic variation, with 27% of variation in inter-individual methylation differences explained by genetic distance. Similar analyses have found significant correlations in reef-building coral (Dimond & Roberts 2020) and humans (Carja et al. 2017). By only focusing on measures of population differentiation, previous marine invertebrate studies may have missed couplings between methylation and genetic patterns. There are three nonexclusive scenarios that could explain the observed relationship between genetic and epigenetic patterns: 1) genetic state results in methylation change (e.g. CpG-SNPs), 2) methylation state results in genetic change, and 3) epigenetic and genetic changes occur in parallel due to independent molecular mechanisms, but are associated through either physical linkage or shared evolutionary pressures (Figure 7).

**Figure 7:**
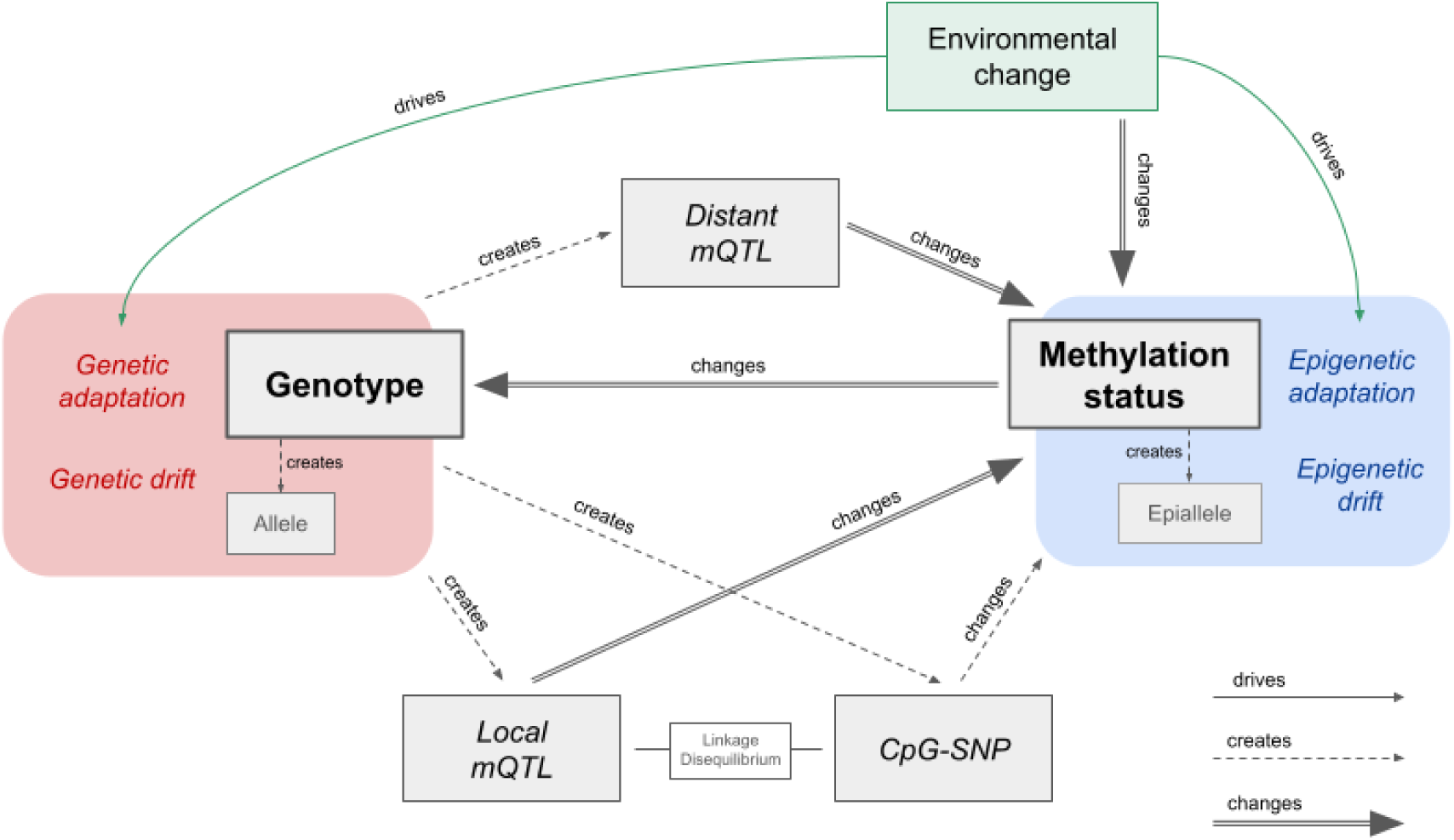
Molecular and evolutionary mechanisms linking genetic variation and methylation variation. DNA mutations can change or create methylation epialleles at CpGs, either directly through CpG-SNPs, or indirectly through the creation of local or distant mQTLs. Some of these mQTL associations will be spurious, due to linkage disequilibrium (LD) with CpG-SNPs or other mQTLs. Methylation epialleles can be created or changed by stochastic epimutations or external signals from the environment. Methylation status in turn can change the rate of DNA mutations at a local scale. Observed epigenetic and genetic associations may instead be due to independent molecular mechanisms that occur in parallel due to shared evolutionary pressures.

CpG-SNPs have been implicated as important drivers of genome-epigenome interactions in vertebrates, either by removing a CpG site on one or both strands and directly disrupting methylation, or by influencing local methylation activity (Zhi et al. 2013; McClay et al. 2015). A considerable proportion of the SNPs in our study were CpG-SNPs (12.3%), 40.1% of which were within 350bp of a methylated CpG, and therefore capable of influencing our MBD-BS measurements. The enrichment of CpG-SNPs associated with methylated CpGs supports the hypothesis that methylation could have preceded and induced genetic variation by altering genome stability and mutation rates (Flores et al. 2013). Methylated cytosines readily mutate to thymine by deamination, which results in an overall depletion of CpG dinucleotides (Coulondre et al. 1978; Schorderet & Gartler 1992; Bird 1980). For instance, in the Pacific oyster *C. gigas* mutation rate is biased towards GC -> AT, particularly at methylated CpG sites, and in coding regions (Song 2020), and genes predicted to have low levels of methylation (analyzed in-silico using the established CpG observed / expected relationship) are less genetically diverse (analyzed via SNPs) (Roberts & Gavery 2012). Alternatively, some CpG-SNPs may have preceded methylation and led to beneficial methylation variation, in which case they may be associated with mQTLs.

High-density methylome and genotyping studies in model taxa have determined that a substantial proportion of variably methylated sites are under local genetic control by mQTLs. To our knowledge, this is the first mQTL analysis in a marine invertebrate. We found 7,166 of tested CpGs were under genetic control, either locally (2.3%) or distantly (3.0%). This is lower than that found for human blood cells (15% local, 0.08% distant)(McClay et al. 2015) and *Arabidopsis thaliana* (18%) (Dubin et al. 2015), although our study has much lower coverage in both methylation and genetic data. The McClay human study also found that 97.7% of SNPs were local meth-QTLs, which is much higher than the 21% found in our study. One likely explanation for this is the highly fragmented status of our draft genome, with 158,535 scaffolds under 50kb in length. It is likely that some SNPs within 50kb of a CpG were actually tested as distant mQTLs. While our mQTL analysis is not entirely comparable to larger scaled studies in humans and plants, it nevertheless shows that associations with genetic variants can be a significant source of variation in methylation, and should therefore be investigated further with whole genome genotyping. CpG-SNPs are one possible mechanism underlying local mQTLs, and we do see an enrichment of CpG-SNPs in local mQTLs compared to distant mQTLs. This result has also been seen in model organisms and humans, however in those cases CpG-SNPs contributed to over 75% of local mQTLs (McClay et al. 2015).

For mQTLs that lack CpG-SNPs, alternative mechanisms must be considered. Binding of transcription factors has been linked to changes in local methylation levels, for example a loss of methylation upon transcription factor binding (Héberlé & Bardet 2019). In this framework, a SNP within a transcription factor binding site may affect methylation locally, while a SNPs that affects the expression or activity of transcription factors could generate changes in methylation wherever the transcription factor binds (Lienert et al. 2011; Martin-Trujillo et al. 2020). Our functional enrichment tests suggest this mechanism may be acting in *O. lurida* by finding genes with mQTL SNPs enriched for “DNA-binding” and “transcription regulation”, and five distant mQTLs SNPs within genes involved in transcription factor complexes. Genetic differences that affect binding of different chromatin classes have also been shown to modulate local methylation patterns (Jeffery & Nakielny 2004; Banovich et al. 2014). One particularly exciting result is that genes containing distantly associated CpGs were highly enriched for RNA processing and binding functions, including multiple RNA binding motif proteins and DEAD-box RNA helicases. DEAD-box RNA helicases are known to co-regulate transcription factors and contribute to chromatin remodeling in multicellular organisms, although the exact molecular mechanisms are still unclear (Giraud et al. 2018). They have also been linked to epigenetic control of abiotic stress-responsive transcription factors in plants through an RNA-directed DNA methylation pathway (Barak et al. 2014). More research integrating chromatin annotations (e.g., ATACseq), CpG methylation, genetic diversity, and gene expression are required to begin elucidating how these mechanisms interact to drive phenotypic divergence.

To confidently state that epigenomic variation is under genetic control for all detected local mQTLs, one assumes that epigenetic inheritance by other means is absent. If epigenetic marks can be inherited between generations, then associations with local genetic variants may simply be due to LD between the segregating epiallele and nearby SNPs. Epigenetic inheritance is well characterized in plants (Taudt et al. 2016), and there is evidence of environmentally-driven epigenetic changes that persist across generations in corals and oysters, although the mechanisms of invertebrate epigenetic inheritance is still not understood (Johnson et al. 2020; Downey-Wall et al. 2020; Lim et al. 2020; Wang et al. 2020; Akcha et al. 2020; Venkataraman et al. 2020). It is also possible that genetic variants and epialleles may be under parallel selection due to phenotype-genotype interactions, which may lead to a spurious mQTL association (Schmid et al. 2018; Taudt et al. 2016). However, since epialleles can undergo both forward and backward changes, epimutation rates are much higher than DNA mutations and therefore spurious mQTL associations will break down rapidly. Comparing mQTL analyses between generations would help identify both the heritability of CpG methylation and the consistency of mQTL results.

### Evolutionary implications

Many marine invertebrates with large ranges experience spatial heterogeneity in abiotic and biotic factors that lead to population-level divergence in fitness-related traits (Sanford & Kelly 2011). This environmentally-driven divergence may be facilitated through phenotypic plasticity, selection for locally-favorable genotypes, or a combination. Here we were able to examine the primary molecular mechanisms underlying plasticity and adaptation: epigenetic modifications and genetic variation. Interestingly, in this system we found a clear coupling of the two, with 27% of individual epigenetic variation attributable to genetics. This result has profound implications for studies of both evolutionary processes and molecular machinery. First, studies of plasticity and epigenetic variation among groups from different environments must also account for genetic variation, rather than attributing all differences to the environment. Second, as genetic variation is clearly heritable, our results suggest that some proportion of DNA methylation (and likely associated phenotypes) are also heritable. Finally, despite our two populations being raised in the same environment, 73% of the epigenetic variation in our system was not attributable to genetics. Characterizing the basis of this additional epigenetic diversity, such as a historical influence of the environment or independent heritable mechanisms, will identify avenues adjacent to genetic adaptation for producing long-term shifts in phenotype.

## Methods

### Draft Genome Assembly and Annotation

To facilitate the analysis of genetic and epigenetic data, a draft genome for the Olympia oyster was developed using a combination of short-read sequence data (Illumina HiSeq4000) combined with long-read sequence data (PacBio RSII) using PBJelly (PBSuite_15.8.24; English et al, 2012). Short reads (NCBI SRA: SRP072461) were assembled using SOAPdenovo (Li et al, 2008). The scaffolds (n=765,755) from this assembly were combined with the PacBio long-read data (NCBI SRA: SRR5809355) using PBJelly (PBSuite_15.8.24; English et al, 2012). Assembly with PBJelly was performed using the default settings. Only contigs longer than 1000 bp were used for further analysis. Genome assembly parameters were compiled using QUAST (v4.5; Gurevich et al, 2013).

Genome annotation was performed using MAKER (v.2.31.10; Campbell et al, 2014) configured to use Message Passing Interface (MPI). A custom repeat library for use in MAKER was generated using RepeatModeler (open-1.0.11; . Hubley and Smit, 2008). RepeatModeler was configured with the following software: RepeatMasker (open-4.0.7; configured with Repbase RepeatMasker v20170127; (Bao et al. 2015), RECON (v1.08; Bao and Eddy, 2002) with RepeatMasker patch, RepeatScout (v1.0.5; Price et al, 2005) and RepeatMaskerBlast (RMBLast (2.6.0)) configured with the isb-2.6.0+-changes-vers2 patch file, and TRF (v4.0.4; Benson, 1999).

MAKER was run on two high performance computing (HPC) nodes (Lenov NextScale, E5-2680 v4 dual CPUs, 28 cores, 128GB RAM) on the University of Washington’s shared scalable compute cluster (Hyak) using the icc_19-ompi_3.1.2 module (Intel C compiler v19, Open MPI v3.1.2). An Olympia oyster transcriptome assembly was used for EST data. Protein data used was a concatenation of NCBI proteomes from *Crassostrea gigas* and *Crassostrea virginica*. *Ab-initio* gene training was performed twice using the included SNAP software (Korf, 2004). Functional protein annotation was performed using BLASTp (v.2.6.0+; Altschul et al, 1990) against a UniProt SwissProt BLAST database (FastA file formatted using BLAST 2.8.1+) downloaded on 01/09/2019. The MAKER functions ‘maker_functional_gff’ and ‘maker_functional_fastà both used the same UniProt SwissProt BLAST database. Protein domain annotation was performed using InterProScan 5 (v5.31-70.0; Jones et al, 2014). Code and data files used for genome annotation are available in the accompanying repository https://github.com/sr320/paper-oly-mbdbs-gen.

### Experimental Design

DNA was extracted from adductor muscle tissue from 184 individuals (88 from Hood Canal and 96 from Oyster Bay), using E.Z.N.A. Mollusc Kit with RNase A treatment (Omega) according to the manufacturer’s instructions. DNA quality was examined on a 1% TAE agarose gel and DNA concentration was determined using the dsDNA BR Assay Kit on a Qubit 2 fluorometer (Invitrogen).

### Genetic Analysis

#### 2bRAD Sequencing and Genotyping

Using a 2b-RAD reduced-representation sequencing approach (Wang et al. 2012), we sequenced 184 individuals and 53 technical replicates from the two Puget Sound populations for a total of 237 samples across 4 lanes of 50bp single-end Illumina HiSeq2500 and 1 HiSeq4000 lane. The frequent-cutter restriction enzyme AlfI was used with modified adaptors (5’-NNR-3’) to target **¼** of all AlfI restriction sites in the genome. We followed the 2bRAD library protocol developed by Eli Meyer (available at https://github.com/sr320/paper-oly-mbdbs-gen), except that we used 900 ng of starting DNA, 19 PCR cycles as determined by a test PCR, and we concentrated the final pooled libraries using a Qiagen PCR kit prior to sequencing. Sequencing and sample demultiplexing was performed by GENEWIZ for the four HiSeq2500 lanes and the University of Chicago’s Functional Genomics Center for the one HiSeq4000 lane. Sequencing of some of these samples was previously described in (Silliman et al. 2018).

Scripts by Mikhail Matz were used for quality filtering, read trimming, and mapping to the reference genome (https://github.com/z0on/2bRAD_denovo). Read trimming was performed by cutadapt (Martin 2011). Samples were retained for mapping to the genome and genotyping if they had greater than 1.3 million reads after filtering. Samples were mapped to the genome using Bowtie2 with the --local option (Langmead & Salzberg 2012). Genotype likelihoods were calculated using ANGSD (Korneliussen et al. 2014) with the following filters: no triallelic sites, p-value that SNP is true 1e-3, minimal mapping quality 20, minimal base quality 25, minimal number of genotyped individuals 80 (∼70% of individuals passing filter), minimal number of reads at a site 3, minimum p-value for strand bias 1e-5, and minimum overall allele frequency 0.01. This filtering retained 114 samples and 5,269 SNPs.

#### Genetic distance, PCA, Admixture

The genotype likelihoods produced by ANGSD were used for examining population genetic structure and estimating pairwise genetic distance. NGSadmix was used to perform an ADMIXTURE analysis based on genotype likelihoods of 3,724 SNPs, after filtering further for a minimum overall allele frequency of 0.05 (Skotte et al. 2013). The most likely number of genetic clusters (K) was determined using the (Evanno et al. 2005) method by running NGSadmix 10 times for each value of K, with K ranging from one to five, and then uploading the results to Clumpak (Kopelman et al. 2015). The q values for the best K were plotted in R. Pairwise genetic distances between all individuals were estimated using ngsDist with default parameters (Vieira et al. 2016). A matrix of genetic distances for the MBD18 samples was subsetted and used for comparative analyses with methylation data.

For a Principal Components Analysis (PCA) of all samples, we used ANGSD to estimate a covariance matrix by sampling a single read at each polymorphic site using the same filtering parameters as previously described. We then performed an eigenvalue decomposition on the matrix and plotted the PCA in R. For the PCA on only the MBD18 samples, we subsetted the covariance matrix and ran an eigenvalue decomposition on those samples alone.

To detect SNPs under putative directional selection, we used qctool v2.0 (https://www.well.ox.ac.uk/~gav/qctool_v2/) to convert our genotype likelihoods to a VCF of SNPs with > 90% confidence. SNPs with less than 90% confidence were coded as missing. We used BayeScan v2.1 (Foll & Gaggiotti 2008) with 1:10 prior odds, 100,000 iterations, a burn-in length of 50,000, a false discovery rate (FDR) of 10%, and default parameters. Results were visualized in R.

To measure population genetic differentiation (F_ST_), we used the realSFS command in ANGSD to estimate the Site Frequency Spectrum (SFS) separately for each population, then calculated the 2D-SFS which was used as a prior for estimating the joint allele frequency probabilities at each site. In order to avoid distorting the allele frequency spectrum, we did not filter our data based on the p-value that a SNP was true or for minimum allele frequency. We then filtered out potential lumped paralogs sites by removing sites where heterozygotes likely compromised more than 75% of all genotypes. This filtering strategy resulted in 363,405 sites and 5,882 SNPs. Global F_ST_ between populations and per-site F_ST_ was calculated using ANGSD, based on (Reynolds et al. 1983). A weighted F_ST_ estimate was calculated for each gene by including all SNPs within ± 2kb of an annotated gene region.

### DNA Methylation

#### MBD-BS Library Preparation and Alignment

DNA was isolated from adductor tissue using the E.Z.N.A. Mollusc Kit (Omega) according to the manufacturer’s protocol. A total of 18 samples were extracted for DNA methylation analysis, 9 from the Hood Canal population and 9 from the Oyster Bay population. Samples were sheared to a target size of 350bp using a Bioruptor 300 (Diagenode) sonicator. Fragmentation was confirmed with a Bioanalyzer 2100 (Agilent). Methylated DNA was selected using the MethylMiner Methylated DNA Enrichment Kit (Invitrogen) according to the manufacturer’s instructions for a single, high-salt elution. Samples were sent to ZymoResearch for bisulfite conversion, and Illumina library preparation for 50bp single-end reads and sequencing with the Pico Methyl-Seq Library Prep Kit (ZymoResearch). Samples were multiplexed into a single library and sequenced on an Illumina HiSeq2500 (Illumina). This library was sequenced across three lanes to achieve the desired number of reads.

Sequence quality was checked by FastQC v0.11.8 and adapters were trimmed using TrimGalore! version 0.4.5 (Andrews 2010; Krueger 2012). Bisulfite-converted genomes were created in-silico with Bowtie 2-2.3.4 (Linux x84_64 version; (Langmead & Salzberg 2012) using bismark_genome_preparation through Bismark v0.21.0 (Krueger & Andrews 2011). Trimmed reads were aligned to these genomes with Bismark v0.21.0. Alignment files were deduplicated with deduplicate_bismark and sorted using SAMtools v.1.9 (Li et al. 2009). Methylation calls were extracted from sorted deduplicated alignment files using coverage2cystosine with -- merge_CpG parameter.

#### General DNA methylation landscape

To assess general methylation patterns in *O. lurida*, quality trimmed MBD-BS reads from all samples (n=18) were concatenated, then re-aligned to the genome using Bismark with settings as described above. Only loci with at least 5x coverage were examined. A cytosine locus was deemed methylated if 50% or more of the reads remained cytosines after bisulfite conversion (Gavery & Roberts 2013; Venkataraman et al. 2020). To characterize methylation landscape, loci were intersected with the following *O. lurida* genome features using bedtools v2.29.0: exons, introns, gene flanking regions (2kb upstream and downstream), transposable elements, and unknown regions (Quinlan 2014). All CpG loci in the *O. lurida* draft genome were similarly annotated to characterize the distribution of candidate CpG methylation sites across features. Using chi-squared contingency tests in R, we examined whether the distribution of methylated loci across genomic features differed from the distribution of all CpG sites in the genome (ɑ=0.05).

#### Comparative methylation analyses

Associations between *O. lurida* population (Hood Canal, South Puget Sound) and methylation patterns were examined by assessing differentially methylated loci (DMLs) and differentially methylated gene regions (DMGs). Bismark alignment files (.bam format) were first processed in methylKit (version 1.8.1) (Akalin et al. 2012) by using `processBismarkAln` to convert them to a `methylRawList` object, which contains per-base methylation calls for each sample. Loci were filtered to retain those with at minimum 5x coverage using `filterByCoverage`, and `unite` selected only loci that were retained across 7 of the 9 samples within each population (N=18). Additional loci were included in the comparative analyses to incorporate loci that were very likely unmethylated in one population but highly methylated in the other, which is not captured in MBDSeq data due to the heavy bias for methylated regions. This was accomplished by identifying CpG loci that were widely sequenced in one population (data present for seven of the nine samples) and minimally sequenced in the other population (data present for one sample or less), and assuming that the samples with no data in the low-sequenced population were unmethylated at 5x coverage. Global differences in methylation patterns were assessed by Principal Component Analysis (PCA) using the `PCASamples` function (a version of `prcomp`), from a percent methylation matrix that was built using `percMethylation`. A matrix of sample x sample manhattan distances was generated from the percent methylation matrix using dist() from the stats package for R v4.0.4 and used for comparative analyses with genetic data.

#### Differentially Methylated Loci (DMLs)

DMLs were determined for each CpG locus using logistic regression in MethylKit with `calculateDiffMeth`, and P-values were adjusted to Q-values using the SLIM method (Wang et al. 2011). Loci with Q-value<0.01 and percent methylation difference >25% were determined to be differentially methylated (DMLs).

#### Differentially Methylated Gene Regions (DMGs)

Gene regions were assessed for differential methylation among populations. Methylated loci that overlapped with known gene regions were identified using the BEDtools `intersectBed` function, a list of known genes that were identified using the genome annotation tool MAKER (Cantarel et al. 2008), and expanded to include 2kb upstream and downstream of gene bodies using BEDtools `slopBed.` Gene regions were assessed individually for differential methylation between oyster populations using binomial GLMs and Chi-square tests (Liew et al. 2018). P-values were adjusted using the Benjamini and Hochberg method (Benjamini & Hochberg 1995). Gene regions that contained fewer than 5 methylated loci were discarded prior to GLM analysis. Epigenetic divergence was estimated by P_ST_ (Johnson & Kelly 2020) for 14,088 random 10kb bins using `Pst` from the Pstat R package (Blondeau Da Silva Stephane [aut 2017).

#### Gene Enrichment Analyses

DMGs and genes that contain DMLs were each tested for enriched biological functions. For each gene set, gene sequences were merged with the *O. lurida* genome to generate a list of Uniprot IDs from annotated genes. Enriched biological processes in each gene set were identified with the Gene-Enrichment and Functional Annotation Tool from DAVID v6.8 as those with modified Fisher Exact P-Values (EASE Scores) <0.1 (Huang et al. 2009).

### Comparing DNA Methylation and Genetics

To investigate the relationship between genetic and DNA methylation variation, we compared summary statistics at both the level of the individual and genomic region for the 18 individuals where we had both genetic and epigenetic data (MBD18). First we compared pairwise genetic distances based on 5,269 SNPs against pairwise Manhattan distances based on all filtered methylation data, and determined both the Pearson and Spearman correlations in R. We also compared the distances when only using DMLs for methylation distances (Figure 4). We then assessed the correlation between the 1st PC scores from SNP data against the 2nd PC scores of methylation data (Figure 3). We also calculated mean F_ST_ and P_ST_ for the 827 10kb genomic bins where we had both SNP and methylation data. These F_ST_ and P_ST_ values were calculated as previously described for gene regions, with overlapping 10kb regions identified with BEDtools (Quinlan 2014). To identify CpG-SNPs in our set of 5,269 SNPs, we used the `injectSNPsMAF` and `getCpGsetCG` functions in the R package RaMWAS v1.18 (Shabalin et al. 2018) and the package bedR v1.0.7 (Haider et al. 2016).

To determine the relationship between regions of the genome with genetic variation and regions with inter-individual methylation variation, we conducted a mQTL analysis using a linear regression model ‘modelLINEAR’ in the R package MatrixEQTL (Shabalin 2012). CpGs were removed if no samples had greater than 12% difference in methylation, resulting in 232,567 CpGs for the analysis. Methylation values were corrected using the inverse quantile normal transformation of ranked values using custom R code (McCaw et al. 2020). 2,860 SNPs remained after filtering for those genotyped in at least 7 individuals of both populations and with an overall MAF > 0.05. To control for ancestry, the first three PCs of the SNP data were included as covariates in the regression model. Local mQTLs were determined to be SNPs within 50kb of the CpG and a p-value threshold of 0.01, while disant mQTLs were greater than 50kb from the CpG or on a different scaffold, had a p-value threshold of 0.01, and an FDR of 1% after Benjamini–Hochberg correction. Summary and plotting of mQTL loci was performed in R and ggplot2 (Wickham 2016). Gene regions containing mQTL SNPs and their associated CpGs were analyzed for functional enrichment with DAVID as described for DMLs, however for the CpGs associated with distant mQTLs we used an EASE score cutoff of 0.05.

## Data Availability Statement

Code and intermediate analysis files used in this study are available in the accompanying repository https://github.com/sr320/paper-oly-mbdbs-gen. The genome assembly can be found at NCBI under the accession PRJEB39287. The annotation files used for these analyses and raw data for the genome assembly will be made available upon publication. Raw 2b-RAD data and MBD-BS data will be available on NCBI Sequence Read Archive by time of submission.

