## Supplementary for "Epigenetic and genetic population structure is coupled in a marine invertebrate"

**1. General *O. lurida* methylation characteristics**

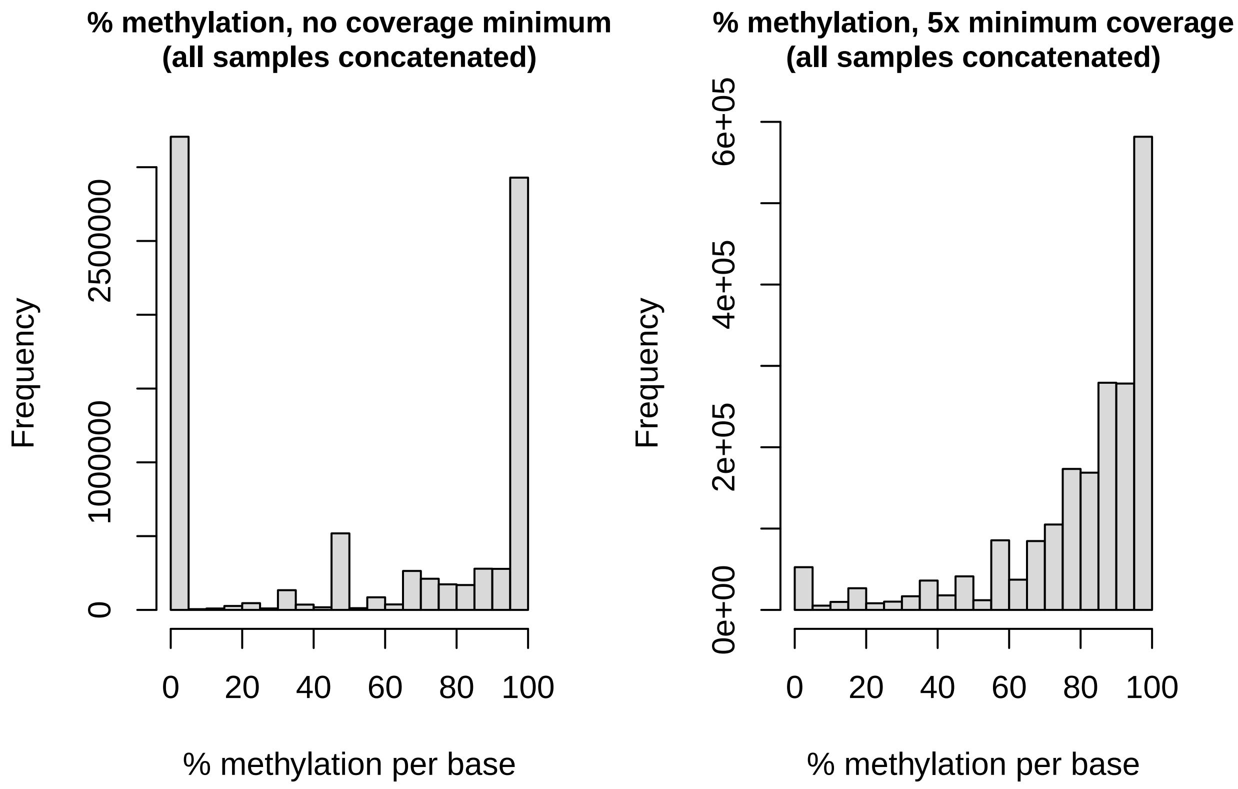


**Supplemental Figure 1**: Frequency distribution of % methylation in *O. lurida* prior to filtering (left) and after filtering for loci with at minimum 5x coverage (right).

| Feature | % of all CpGs in genome | % of all methylated loci | Ratio of % meth:%CpG |
|---|---|---|---|
| Exon | 3.99% | 14.71% | 3.69 |
| Intron | 13.94% | 19.76% | 1.42 |
| 5' flanking region (-2kb) | 3.91% | 4.64% | 1.19 |
| 3' flanking region (+2kb) | 3.92% | 4.66% | 1.19 |
| Transposable Elements | 14.93% | 13.83% | 0.93 |
| Unknown genome regions | 56.01% | 32.25% | 0.58 |

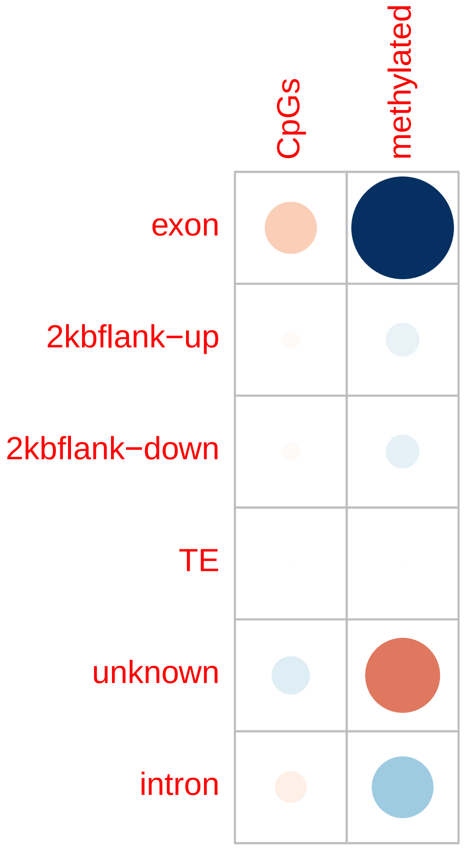


**Supplemental Figure 2**: Top: Table showing the % of all CpG loci and methylated loci that overlap with each genomic feature. Bottom: A correlation plot, showing residuals from chi-squared tests of homogeneity on the distribution of methylated loci and CpG loci that intersect with features. Blue=positive associations, red=negative associations, and size of circles represents the absolute value of each correlation coefficient.

---

---

**Supplemental Table 1**: Biological functions that are enriched in the *Ostrea lurida* genome. Also available in .csv format: https://github.com/sr320/paper-oly-mbdbs-gen/blob/master/analyses/methylation-characteristics/methylated-loci-enriched-BP.csv

| GO Term | Biological Process | PValue | FDR | GO slim |
|---|---|---|---|---|
| GO:0007067 | mitotic nuclear division | 0 | 0.37 | cell cycle and proliferation |
| GO:0007049 | cell cycle | 0 | 0.66 | cell cycle and proliferation |
| GO:0051321 | meiotic cell cycle | 0.05 | 1 | cell cycle and proliferation |
| GO:0000082 | G1/S transition of mitotic cell cycle | 0.07 | 1 | cell cycle and proliferation |
| GO:0000086 | G2/M transition of mitotic cell cycle | 0.09 | 1 | cell cycle and proliferation |
| GO:0007067 | mitotic nuclear division | 0 | 0.37 | cell organization and biogenesis |

| GO Term | Biological Process | PValue | FDR | GO slim |
|---|---|---|---|---|
| GO:0016569 | covalent chromatin modification | 0.01 | 1 | cell organization and biogenesis |
| GO:0060271 | cilium morphogenesis | 0.01 | 1 | cell organization and biogenesis |
| GO:0042384 | cilium assembly | 0.01 | 1 | cell organization and biogenesis |
| GO:0007030 | Golgi organization | 0.02 | 1 | cell organization and biogenesis |
| GO:0030030 | cell projection organization | 0.05 | 1 | cell organization and biogenesis |
| GO:0007005 | mitochondrion organization | 0.05 | 1 | cell organization and biogenesis |
| GO:0001822 | kidney development | 0.02 | 1 | developmental processes |
| GO:0001843 | neural tube closure | 0.07 | 1 | developmental processes |
| GO:0006281 | DNA repair | 0 | 0.08 | DNA metabolism |
| GO:0006260 | DNA replication | 0 | 0.37 | DNA metabolism |
| GO:0006310 | DNA recombination | 0.04 | 1 | DNA metabolism |
| GO:0006302 | double-strand break repair | 0.08 | 1 | DNA metabolism |
| GO:0006284 | base-excision repair | 0.09 | 1 | DNA metabolism |
| GO:0051301 | cell division | 0 | 0.12 | other biological processes |
| GO:0007018 | microtubule-based movement | 0.02 | 1 | other biological processes |
| GO:0042254 | ribosome biogenesis | 0.05 | 1 | other biological processes |
| GO:0016032 | viral process | 0.05 | 1 | other biological processes |
| GO:0043547 | positive regulation of GTPase activity | 0.05 | 1 | other biological processes |
| GO:0000910 | cytokinesis | 0.06 | 1 | other biological processes |
| GO:0032092 | positive regulation of protein binding | 0.08 | 1 | other biological processes |
| GO:0008104 | protein localization | 0.09 | 1 | other biological processes |
| GO:0031047 | gene silencing by RNA | 0.01 | 1 | other metabolic processes |
| GO:0006511 | ubiquitin-dependent protein catabolic process | 0 | 0.12 | protein metabolism |
| GO:0006412 | translation | 0 | 0.37 | protein metabolism |
| GO:0050821 | protein stabilization | 0.01 | 1 | protein metabolism |
| GO:0006468 | protein phosphorylation | 0.01 | 1 | protein metabolism |

| GO Term | Biological Process | PValue | FDR | GO slim |
| --- | --- | --- | --- | --- |
| GO:0006413 | translational initiation | 0.03 | 1 | protein metabolism |
| GO:0000209 | protein polyubiquitination | 0.06 | 1 | protein metabolism |
| GO:0042787 | protein ubiquitination involved in ubiquitin-dependent protein catabolic process | 0.08 | 1 | protein metabolism |
| GO:0016567 | protein ubiquitination | 0.09 | 1 | protein metabolism |
| GO:0000398 | mRNA splicing, via spliceosome | 0 | 0.53 | RNA metabolism |
| GO:0006397 | mRNA processing | 0 | 1 | RNA metabolism |
| GO:0006364 | rRNA processing | 0.01 | 1 | RNA metabolism |
| GO:0006396 | RNA processing | 0.01 | 1 | RNA metabolism |
| GO:0008380 | RNA splicing | 0.02 | 1 | RNA metabolism |
| GO:0006366 | transcription from RNA polymerase II promoter | 0.05 | 1 | RNA metabolism |
| GO:0008033 | tRNA processing | 0.06 | 1 | RNA metabolism |
| GO:0006355 | regulation of transcription, DNA-templated | 0.09 | 1 | RNA metabolism |
| GO:0006974 | cellular response to DNA damage stimulus | 0 | 0.05 | stress response |
| GO:0006281 | DNA repair | 0 | 0.08 | stress response |
| GO:0006302 | double-strand break repair | 0.08 | 1 | stress response |
| GO:0006284 | base-excision repair | 0.09 | 1 | stress response |
| GO:0015031 | protein transport | 0 | 0.23 | transport |
| GO:0006886 | intracellular protein transport | 0 | 0.81 | transport |
| GO:0006888 | ER to Golgi vesicle-mediated transport | 0 | 0.96 | transport |
| GO:0016192 | vesicle-mediated transport | 0.02 | 1 | transport |
| GO:0006406 | mRNA export from nucleus | 0.04 | 1 | transport |
| GO:0006606 | protein import into nucleus | 0.09 | 1 | transport |
| GO:0042147 | retrograde transport, endosome to Golgi | 0.1 | 1 | transport |
| GO:0098609 | cell-cell adhesion | 0.04 | 1 | NA |
| GO:0070936 | protein K48-linked ubiquitination | 0.07 | 1 | NA |
| GO:0003341 | cilium movement | 0.07 | 1 | NA |

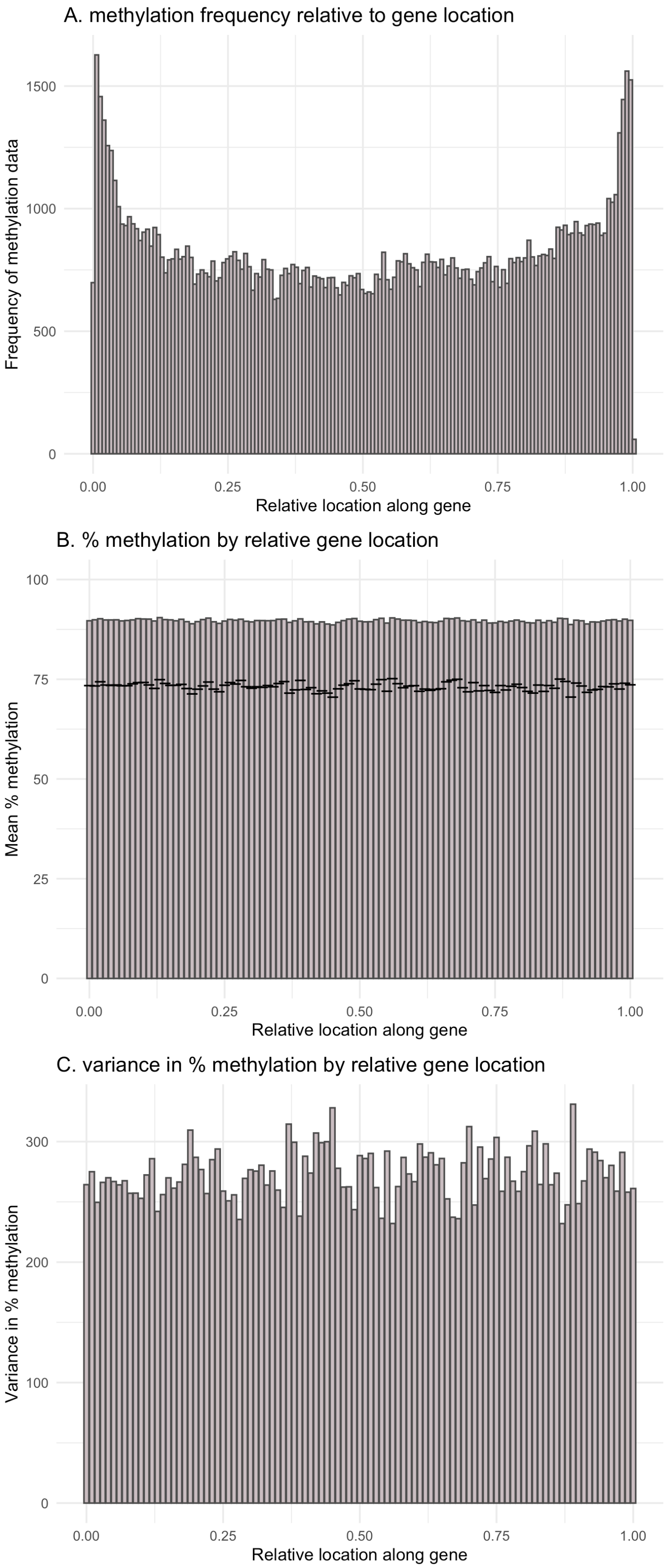


Figure 1: **Supplemental Figure 3:** Gene body methylation patterns. A) relative location of methylation within gene bodies in *Ostrea lurida*. For this analysis methylated loci were designated as those with >50% methylation when averaged across all MBD18 samples, B) percent methylation across a gene body averaged across all MBD18 samples, and C) variation in percent methylation across a gene body, across all MBD18 samples.

**2. Differential methylation analysis among populations**

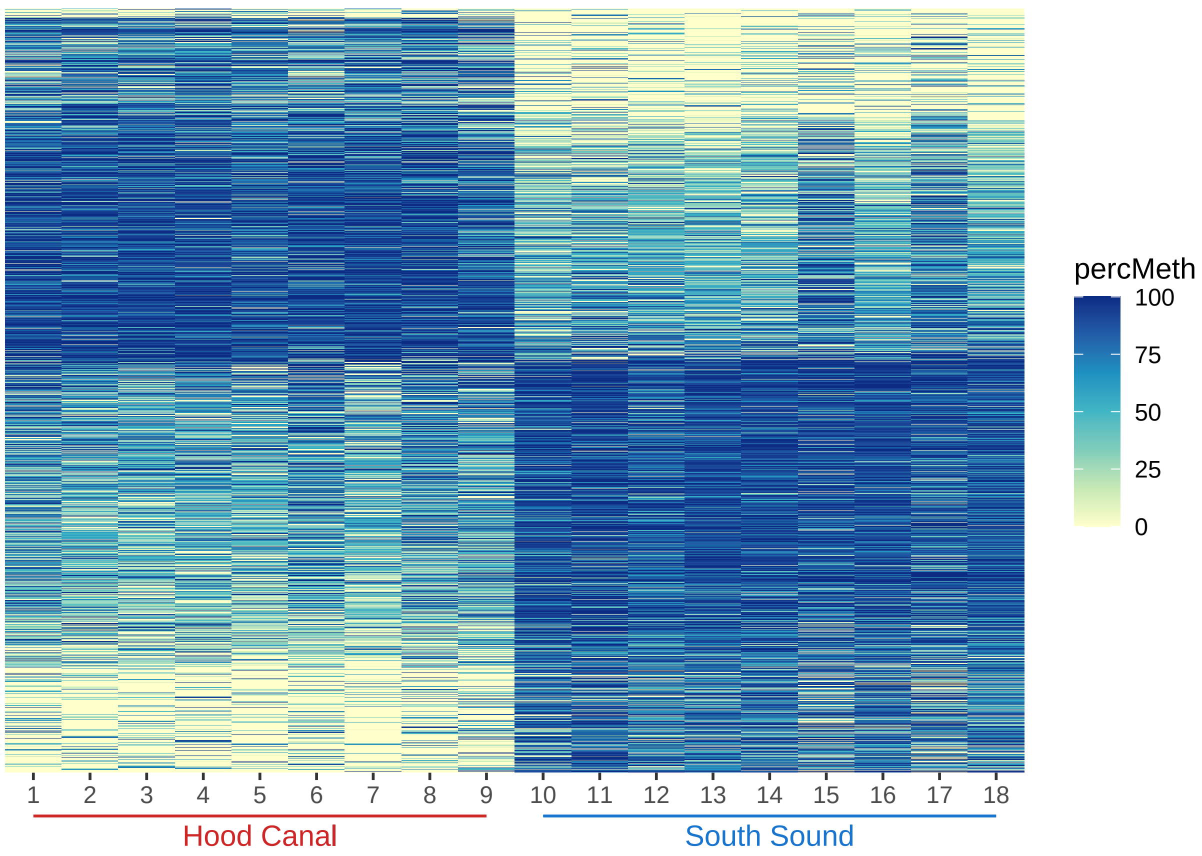


**Supplemental Figure 4**: Heat map of loci that are differentially methylated (DMLs) between the two populations.

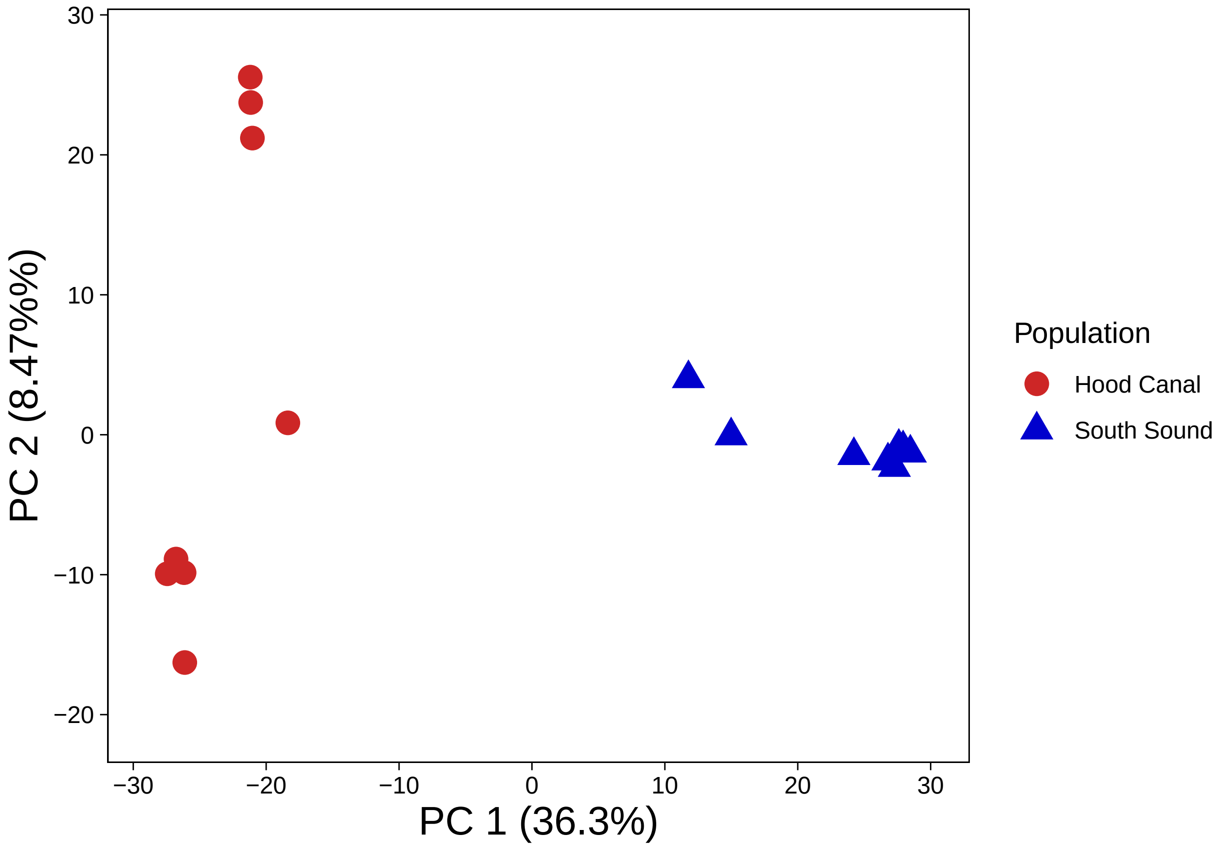


**Supplemental Figure 5**: PCA of methylation data using DMLs only

---

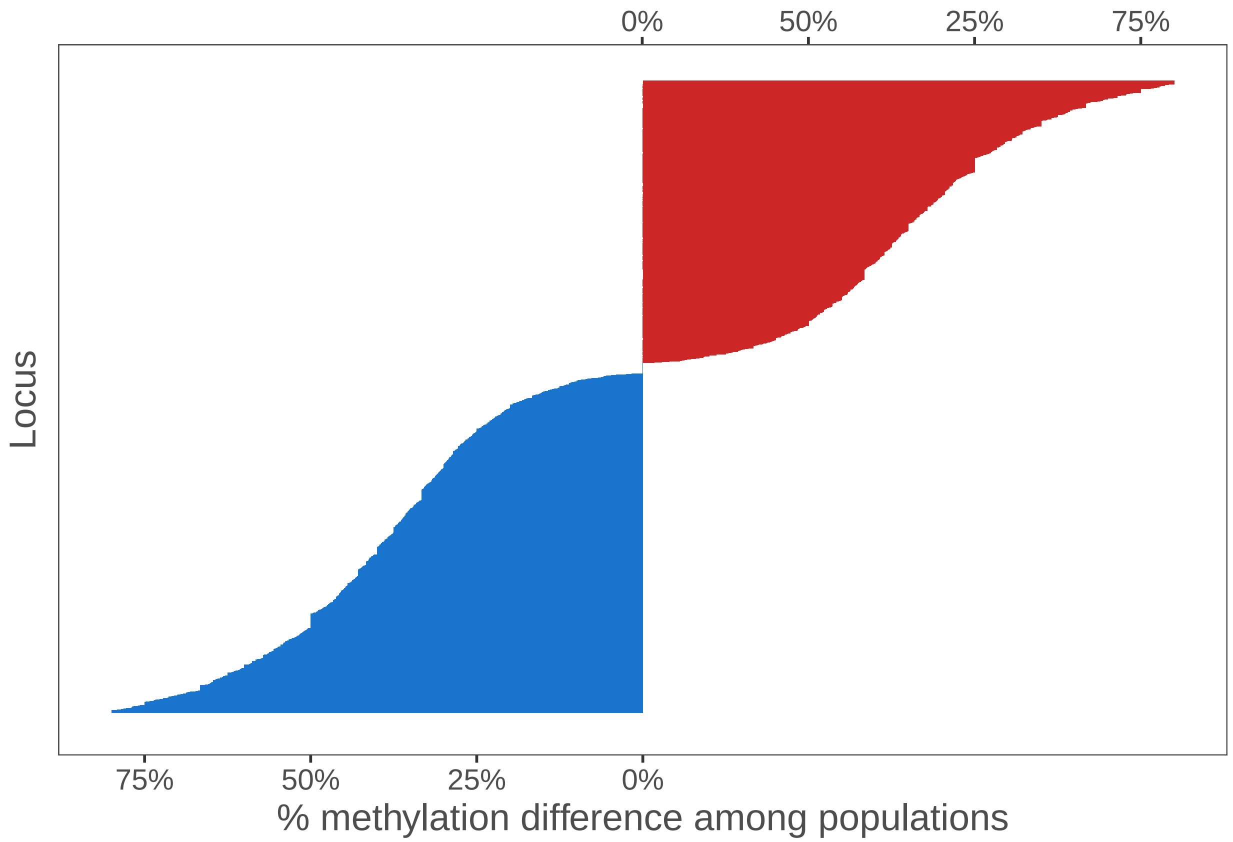


**Supplemental Figure 6**: The % difference between each population's mean methylation for differentially methylated loci, which highlights that both populations have a similar number of hyper/hypo-methylated loci. Loci are sorted by % methylation difference.

---

**Supplemental Table 2**: GO terms and GO Slim terms of biological functions enriched in DMGs & DMLs. EASE scores shown. FDR for all terms is 1.0. Also available in .csv format: https://github.com/sr320/paper-oly-mbdbs-gen/blob/master/analyses/DMG-DML-EnrichedBP.csv

| Go Slim | Term | Function | DMG | DML |
|---|---|---|---|---|
| cell adhesion | GO:0007155 | cell adhesion | 0.012 | |
| cell adhesion | GO:0007156 | homophilic cell adhesion via plasma membrane adhesion molecules | 0.026 | 0.00091 |
| cell adhesion | GO:0016339 | calcium-dependent cell-cell adhesion via plasma membrane cell adhesion molecules | | 0.083 |
| cell organization and biogenesis | GO:0007015 | actin filament organization | 0.014 | |
| cell organization and biogenesis | GO:0048675 | axon extension | 0.047 | |
| cell organization and biogenesis | GO:0007411 | axon guidance | 0.063 | |

| Go Slim | Term | Function | DMG | DML |
|---|---|---|---|---|
| cell organization and biogenesis | GO:0000904 | cell morphogenesis involved in differentiation | 0.095 | |
| cell organization and biogenesis | GO:0030198 | extracellular matrix organization | 0.033 | |
| cell organization and biogenesis | GO:0032836 | glomerular basement membrane development | 0.046 | |
| cell organization and biogenesis | GO:0007040 | lysosome organization | 0.047 | |
| cell organization and biogenesis | GO:0031023 | microtubule organizing center organization | 0.095 | |
| cell organization and biogenesis | GO:0051259 | protein oligomerization | 0.098 | |
| cell organization and biogenesis | GO:0048841 | regulation of axon extension involved in axon guidance | 0.033 | |
| cell organization and biogenesis | GO:0045214 | sarcomere organization | 0.069 | 0.019 |
| cell organization and biogenesis | GO:0055003 | cardiac myofibril assembly | | 0.082 |
| cell organization and biogenesis | GO:0032438 | melanosome organization | | 0.021 |
| cell organization and biogenesis | GO:0045332 | phospholipid translocation | | 0.0058 |
| death | GO:0008625 | extrinsic apoptotic signaling pathway via death domain receptors | 0.095 | |
| developmental processes | GO:0048675 | axon extension | 0.047 | |
| developmental processes | GO:0007411 | axon guidance | 0.063 | |
| developmental processes | GO:0032836 | glomerular basement membrane development | 0.046 | |
| developmental processes | GO:0048841 | regulation of axon extension involved in axon guidance | 0.033 | |
| developmental processes | GO:0045214 | sarcomere organization | 0.069 | 0.019 |
| developmental processes | GO:0007423 | sensory organ development | 0.095 | |
| developmental processes | GO:0055003 | cardiac myofibril assembly | | 0.082 |
| developmental processes | GO:0060429 | epithelium development | | 0.071 |

| Go Slim | Term | Function | DMG | DML |
|---|---|---|---|---|
| developmental processes | GO:0040027 | negative regulation of vulval development | | 0.082 |
| developmental processes | GO:0021942 | radial glia guided migration of Purkinje cell | | 0.082 |
| developmental processes | GO:0050767 | regulation of neurogenesis | | 0.0058 |
| developmental processes | GO:0060438 | trachea development | | 0.082 |
| developmental processes | GO:0001570 | vasculogenesis | | 0.047 |
| other biological processes | GO:0016477 | cell migration | 0.029 | 0.032 |
| other biological processes | GO:0007281 | germ cell development | 0.048 | |
| other biological processes | GO:0019915 | lipid storage | 0.043 | |
| other biological processes | GO:0048477 | oogenesis | 0.046 | |
| other biological processes | GO:0040008 | regulation of growth | 0.032 | |
| other biological processes | GO:0042254 | ribosome biogenesis | 0.092 | |
| other biological processes | GO:0019233 | sensory perception of pain | 0.016 | |
| other biological processes | GO:0006879 | cellular iron ion homeostasis | | 0.047 |
| other biological processes | GO:0050982 | detection of mechanical stimulus | | 0.082 |
| other biological processes | GO:0050801 | ion homeostasis | | 0.082 |
| other biological processes | GO:0007017 | microtubule-based process | | 0.082 |
| other biological processes | GO:0032465 | regulation of cytokinesis | | 0.057 |
| other biological processes | GO:0006941 | striated muscle contraction | | 0.082 |
| other metabolic processes | GO:0042157 | lipoprotein metabolic process | 0.095 | |
| other metabolic processes | GO:0010508 | positive regulation of autophagy | 0.063 | |
| other metabolic processes | GO:0010923 | negative regulation of phosphatase activity | | 0.028 |
| protein metabolism | GO:0031398 | positive regulation of protein ubiquitition | 0.098 | 0.093 |
| protein metabolism | GO:0016567 | protein ubiquitition | | 0.0033 |

| Go Slim | Term | Function | DMG | DML |
|---|---|---|---|---|
| protein metabolism | GO:0042787 | protein ubiquitition involved in ubiquitin-dependent protein catabolic process | | 0.003 |
| signal transduction | GO:0046426 | negative regulation of JAK-STAT cascade | | 0.071 |
| stress response | GO:0042594 | response to starvation | | 0.03 |
| transport | GO:0006895 | Golgi to endosome transport | 0.043 | 0.019 |
| transport | GO:0034220 | ion transmembrane transport | 0.086 | 0.00049 |
| transport | GO:0008333 | endosome to lysosome transport | | 0.083 |
| transport | GO:0045332 | phospholipid translocation | | 0.0058 |
| NA | GO:0044331 | cell-cell adhesion mediated by cadherin | 0.033 | 0.071 |
| NA | GO:0072015 | glomerular visceral epithelial cell development | 0.095 | |
| NA | GO:0086010 | membrane depolarization during action potential | 0.095 | |
| NA | GO:1903955 | positive regulation of protein targeting to mitochondrion | 0.0082 | |
| NA | GO:0038061 | NIK/NF-kappaB sigling | | 0.071 |
| NA | GO:0090175 | regulation of establishment of plar polarity | | 0.082 |

#### 3. Genetic Structure

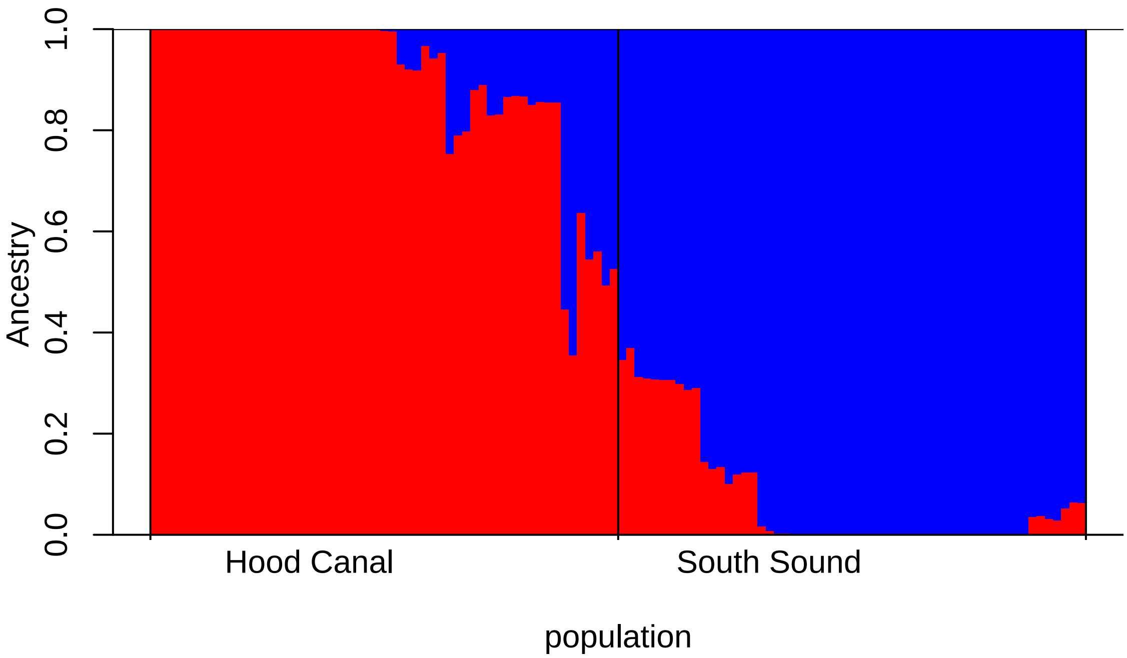


**Supplemental Figure 7**: Admixture plot for 114 individuals at K=2 (determined to be the best K using the Evanno method), based on 3,724 SNPs.

---

**Supplemental Table 3**: Annotations for outlier SNPs that fall within 2kb of a gene. Also available in .bed format, HCSS_Afilt32m70_01_BS-gene2kb.bed

| SNP Contig | SNP Position | Feature Contig | Feature Start | Feature End | Feature Type | Genome ID | Gene Name | GO Terms |
|---|---|---|---|---|---|---|---|---|
| Contig29033 | 27589 | Contig29033 | 11407 | 32648 | gene | OLUR_00003721 | Note=Similar to G2/mitotic-specific cyclin-B (Hydra viridissima OX%3D6082); | Ontology_term=GO:0005634; |
| Contig77382 | 11621 | Contig77382 | 1559 | 12918 | gene | OLUR_00016056 | Note=Similar to Socs5: Suppressor of cytokine signaling 5 (Mus musculus OX%3D10090); | NA |

---

**Supplemental Table 4**: DAVID enrichment results for genes with FST > 0.3. Also available in .tsv format, fst_g03_david.tab

| Category | Term | Count | Percent Genes | PValue | Fold Enrichment | FDR |
|---|---|---|---|---|---|---|
| UP_KEYWORDS | Zinc-finger | 7 | 26.92 | 0.0198 | 2.97 | 1 |
| GOTERM_BP_DIRECT | GO:0006897~endocytosis | 3 | 11.54 | 0.0236 | 11.34 | 1 |
| GOTERM_BP_DIRECT | GO:0006915~apoptotic process | 4 | 15.38 | 0.0281 | 5.50 | 1 |
| UP_SEQ_FEATURE | compositionally biased region:Ser-rich | 4 | 15.38 | 0.0584 | 4.19 | 1 |
| GOTERM_BP_DIRECT | GO:0043401~steroid hormone mediated signaling pathway | 2 | 7.69 | 0.0620 | 30.24 | 1 |
| UP_SEQ_FEATURE | region of interest:Ligand-binding | 2 | 7.69 | 0.0693 | 27.21 | 1 |
| GOTERM_BP_DIRECT | GO:0010506~regulation of autophagy | 2 | 7.69 | 0.0917 | 20.16 | 1 |
| UP_KEYWORDS | Zinc | 7 | 26.92 | 0.0969 | 2.05 | 1 |
| INTERPRO | IPR000536:Nuclear hormone receptor, ligand-binding, core | 2 | 7.69 | 0.0972 | 19.12 | 1 |
| INTERPRO | IPR001628:Zinc finger, nuclear hormone receptor-type | 2 | 7.69 | 0.0972 | 19.12 | 1 |
| INTERPRO | IPR001723:Steroid hormone receptor | 2 | 7.69 | 0.0972 | 19.12 | 1 |

---

### 4. mQTL analysis

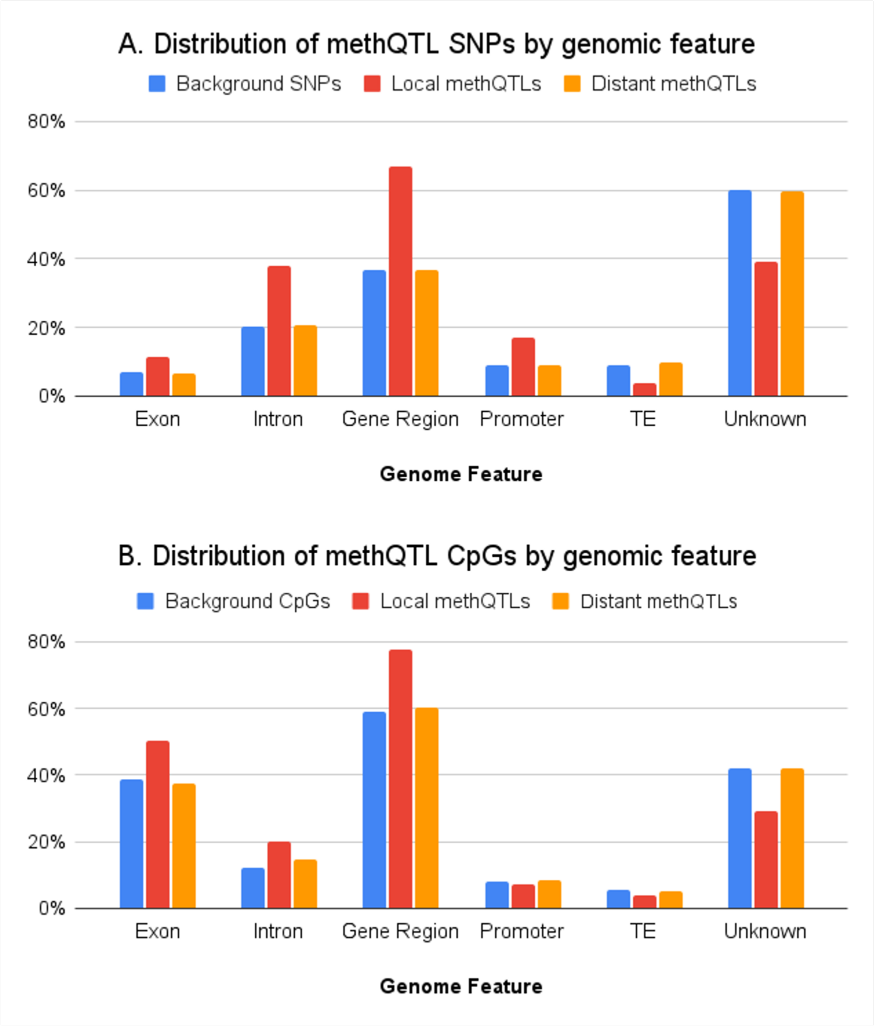


Figure 2: **Supplemental Figure 9:** Comparison of mQTL results that intersect with the following genomic features: exon, intron, promoter region (within 2kb of the 5' end of a gene), gene region (genes plus 2kb upstream and downstream), transposable element, and unknown region of genome. A) SNPs designated as either local (red, within 50kb) or distant (orange) mQTLs, compared with the background (blue) of all SNPs used in the mQTL analysis. B) CpGs associated with either local (red) or distant (orange) mQTLs, compared to the background (blue) of all CpGs used in the mQTL
